# Exploring repeats in rice genomes: Identification, Characterization and its Applications

**DOI:** 10.1101/2022.01.24.477639

**Authors:** Gourab Das, Indira Ghosh

**Affiliations:** Bioinformatics & Computational Biology Facility (BCBF), Advanced Centre for Treatment, Research and Education in Cancer (ACTREC), Tata Memorial Centre (TMC) Sector 22, Utsav Chowk - CISF Rd, Owe Camp, Kharghar, Navi Mumbai, Maharashtra 410210; School of Computational and Integrative Sciences, Jawaharlal Nehru University, New Delhi-110067, India

**Keywords:** Repetitive sequences, Abiotic stress, *Oryza*, Pointwise mutual information, Evolution

## Abstract

Biodiversity is a fundamental property of all natural systems existing in the field biology. It refers to the underlying heterogeneity at different levels of ecology, genetics and evolution. In case of plant systems, dramatic variability has been observed during the Anthropocene at different spatial scales. Environmental stress is one of the major influencing factors behind this plant biodiversity. Huge genetic diversity has been also demonstrated across varieties of important crop species like rice. Repetitive sequences which are a major contributor of genomic diversity in polyploidy plants have been found to occur ubiquitously in their genomes. To date diverse repeat types have been characterized in the plant genomes performing various functions starting from qualitative trait markers to genome evolution and stress management. With an objective to identify of plant stress associated genes using DNA repeat probes, a robust method has been developed. The method has been modularized into three distinct sections. First part is dedicated for identification of different types of repeats. Earlier review has suggested building a pipeline of multiples tools for capturing different types of repeats from the genome sequences. Specialized tools like TROLL, Tandem repeat finder (TRF), PHOBOS and database like REPBASE have been selected for performing this job. Second module is intended for screening of stress related genes from the published articles and databases and the last module has been designed for the association mining between genomic repetitive patterns with stress phenotypes. The method has been used to explore stress associated repeats from 9 *Oryza* species from different continents and other plants like *Arabidopsis* and *Brachypodium*. In case of *Oryza* species distribution repeats has been found to be significantly different between stress associated and housekeeping genes. More than 55% of the repeats are found to be in positive association (nPMI > 0) whereas 26% of the repeats are false-positives. These repetitive probes have been utilized in several applications. Firstly, using as molecular markers to identify stress related genes in different *Oryza* species where availability is limited. Secondly, using as a probe to reanalyzing the evolutionary lineage of *Oryza* species etc.

## Introduction

Rice is one of the mostly used cereals that have been served as a staple food for approximately half of the world population [1]. Along with increasing heat stress due to uprising global warming, several other abiotic stressors including drought, salinity, chemicals, greenhouse gases and biotic stressors like bacteria, fungus viroids pose adverse influence on the growth of the rice plant and its yield [2–6]. Repetitive sequences are abundant in rice genomes [7–10] majority being transposable elements (TEs) [11–13] covering more than 70% of the genome and often play key roles in stress adaptation and disease resistance through many genetic and epigenetic regulatory mechanisms [14–15]. Earlier researches have shown the presence of tandem repeats in excess in defense responsive genes [16]. Microsatellites have become a popular probe for marking candidate genes related to salinity [17], biotic and abiotic stresses [18–21] nowadays. Moreover, influence of transposable elements (TEs) have been significantly observed in genome evolution [22], stress response [23–25], regulations through microRNAs [26], long non-coding RNAs [27] and in many more. In this context, studying repeats in rice genomes to investigate its functional association in abiotic stress management and other applications will be highly demanding and beneficial for better understanding the rice biology.

*Oryza sativa*, a model cultivated species of Asian rice with AA genome, is a diploid angiosperm having one embryonic leaf in their seeds (monocot) and possess 24 chromosomes (2n=24) though one-half of the rice species are allopolyploid [28–29]. In *Oryza* species, total 10 different types of genomes have been characterized which includes 6 diploids (AA, BB, CC, EE, FF, GG) and four allotetraploids (BBCC, CCDD, HHJJ, HHKK) with 48 chromosomes (2n=48) [28]. It is believed that *Oryza* genus has emerged from Gondwanaland approximately 130 million years ago [30] though the debate is still ongoing regarding the actual origin of rice. Both subspecies *O. sativa* ssp. *japonica* (short grain) and *Oryza sativa* ssp. *indica* (long grain) have been domesticated from wild grass ancestor *Oryza rufipogon* around 10000-14000 years ago which is another controversial and live topic of research [31]. *O. rufipogon* (AA genome) is the perennial progenitor of *O. sativa* and wild species *Oryza nivara* (AA genome) is thought to be the recent annual predecessor [32]. African rice *Oryza glaberrima* (AA genome) is believed to be domesticated from wild species *Oryza barthii* (AA genome) around 2000-3000 years ago [33]. Other African wild rice species *Oryza punctata* which possess BB type genome has its own importance for showing resistance to biotic stressors like bacteria, planthoppers etc. Another African wild rice *Oryza brachyantha* is the only known rice species with FF type genome. The only diploid species from Latin America is *Oryza glumaepatula* (AA genome) which has a significant similarity with *O. sativa* genome as estimated by Fuchs et al. [34]. All of these species from different geographical locations with various types of genomes indicates its rapid diversification by dynamic evolution for legitimate adaptation under stressful conditions [35–36]. This provides a suitable platform to perform genome-wide comparative analysis of repetitive elements among rice species exploring its association with several biosynthesis, biogenesis and stress related pathways.

In the present study, different types of TRs and TEs have been identified, characterized and compared among multiple wild type and cultivated genomes to mine their abundance and strength of association with stress related genes. Repeats have been also used as probes for detecting novel stress genes in rice genomes and analyzing evolutionary relationship. These repeats can be served as a knowledgebase for rice stress gene markers.

## Materials and Methods

The complete workflow for the proposed research has been presented in Figure 1. It consists of three modules. A. Identification and characterization of the repetitive sequences from genomes. B. Screening of stress related genes from the published articles and databases and C. Mining association between repeats and stress.

**Figure 1:**
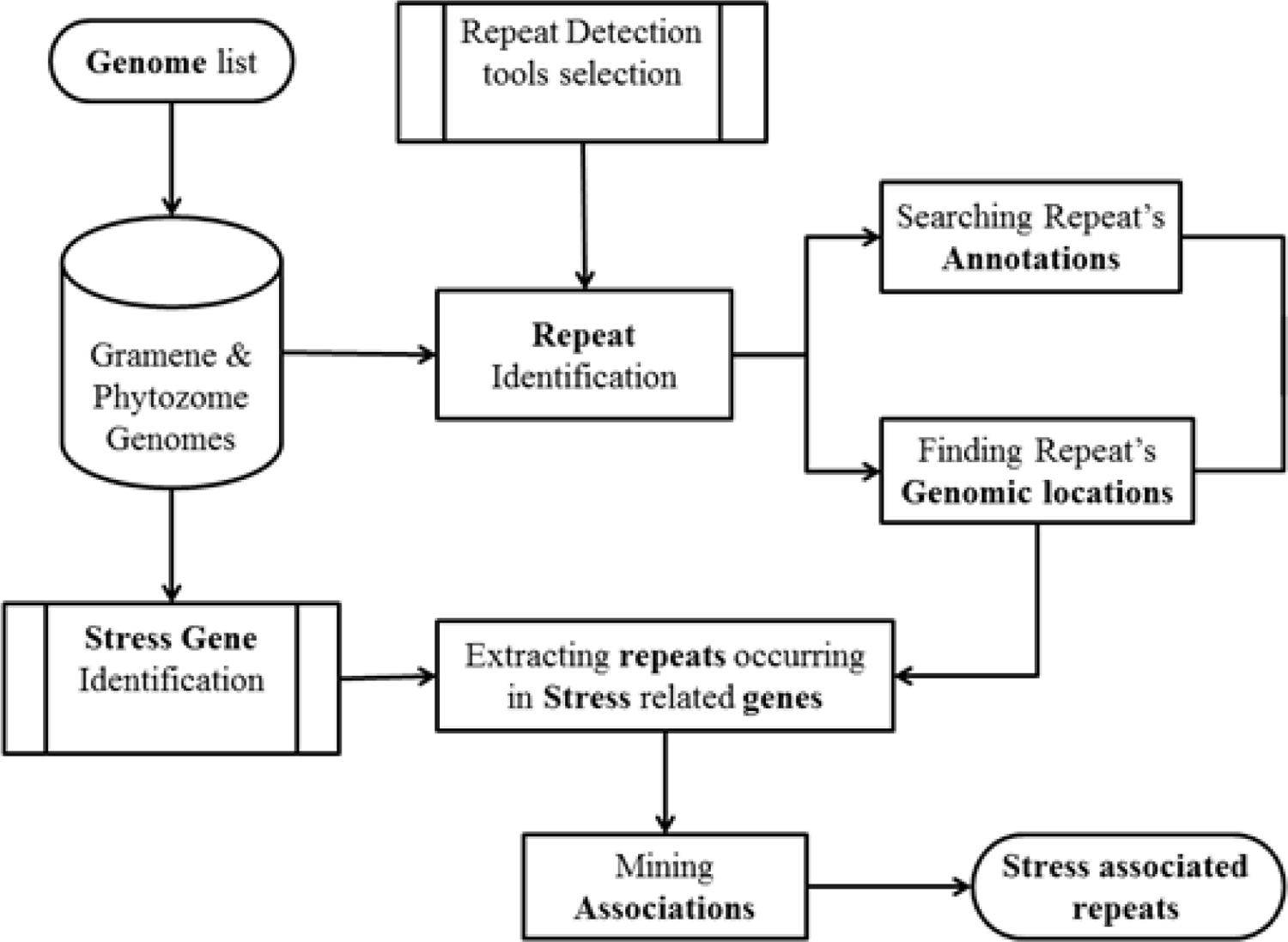
Basic workflow for identification, characterization of repeats in rice genomes and mining its association with stress.

### Genome datasets and annotations

Chromosome sequences of 9 *Oryza* species and other model species *Brachypodium distachyon* (monocot) and *Arabidopsis thaliana* (eudicot) have been downloaded from Gramene-Ensembl database (release 34) [37] using file transfer protocol services (ftp://ftp.gramene.org/pub/gramene/). All protein-coding and non-coding RNA genes available in gff3 format have been parsed for further use.

Major resources of plants complete genomes include Phytozome, NCBI RefSeq, Gramene-Ensembl etc. Phytozome is the Plant Comparative Genomics portal developed by Department of Energy’s Joint Genome Institute (JGI) with genome sequences and annotations of 81 green plant species including the genome of *O. sativa* ssp *japonica* only like NCBI Reference Sequence (RefSeq) database. But Gramene [38] provides the platform for inter-genus comparison with 9 *Oryza* genomes (now 11 in the recent release) so that consistency in results can be checked.

Regarding earlier map-based sequence has provided a rice genome assembly of size 389 Mbp with 95% coverage divided into 12 chromosomes [39] But, few months back a nearly complete de novo assembly of *O. indica* rice genome of estimated size of 390 Mbp and 99% coverage [40]. Table 1 shows the list of *Oryza* genomes used in the present study for comparative analysis.

**Table 1:** List of Rice and other genomes used in the present study

| No. | Common Name | Scientific name | Assembly | Annotation |
| --- | --- | --- | --- | --- |
| 1 | African wild rice | *Oryza barthii* | GCA_000182155.1 | OGE Maker<br>(Aug 2013) |
| 2 | Wild rice from<br>tropical Africa | *Oryza brachyantha* | GCA_000231095.2 /<br>OGEv1.4b | OGEv1.4 |
| 3 | Wild red rice from<br>Africa | *Oryza punctata* | GCA_000573905.1 | OGE Maker<br>(Aug 2013) |
| 4 | African rice | *Oryza glaberrima* | GCA_000147395.1 /<br>AGI1.1 (May 2011) | 2011-05-AGI<br>(MIPS) |
| 5 | Wild rice from<br>America (South) | *Oryza glumaepatula* | GCA_000576495.1 | OGE Maker<br>(Aug 2013) |
| 6 | Wild progenitor of<br>Asian rice | *Oryza rufipogon* | PRJEB4137 | OGE |
| 7 | Wild progenitor of<br>Asian rice | *Oryza nivara* | GCA_000576065.1 | OGE Maker<br>(Aug 2013) |
| 8 | Asian rice | *Oryza sativa* ssp.<br>*japonica* | IRGSP-1.0 | MSU 7.0 |
| 9 | Asian rice | *Oryza sativa* ssp.<br>*indica* | ASM465v1 | 2010-07-BGI |
| **Other model species selected in the present study** |  |  |  |  |
| 1 | Arabidopsis/Thale<br>cress | *Arabidopsis thaliana* | TAIR 9 | TAIR 10 |
| 2 | Stiff Brome grasses | *Brachypodium*<br>*distachyon* | V1.0 | V1.0 |

### Extraction of repeats

In the present study, different types of repetitive patterns have been explored in the rice genomes which include perfect, imperfect types of tandem repeats and many families of transposable elements. Tandem repeats of motif lengths 1-100 bp have been searched in the genomes using selected tools and parameters listed in Table 2. For interspersed repeats like transposable elements, the expert curetted library from Repbase [41] has been used for further research work. Many new repetitive sequence databases have developed recently [42–43] which can be also used to validate the repeat mining.

**Table 2:**
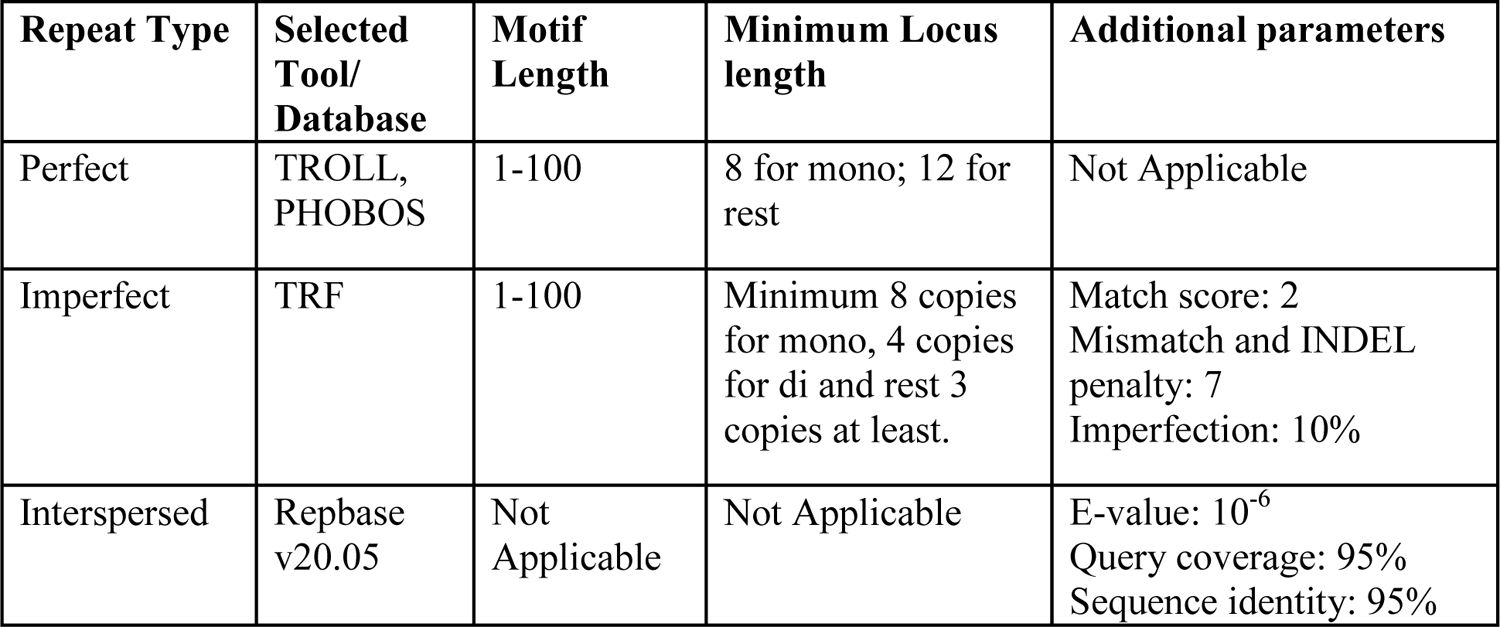
List of parameters for repeat identification

### Tandem Repeats

Tandem repeats have been searched in the genomes using TROLL [44] for perfect microsatellites, PHOBOS [45] for perfect minisatellites and TRF [46] for imperfect repeats. In many of the recent papers, minimum repeat locus length criteria have been used as follows: mono: 8/12, di: 8, tri: 9, tetra: 12, penta: 15, hexa: 18 and for rest of the minisatellites at least two copies of the repeating motif. In the present study, relaxed locus length criteria for mononucleotide repeats have been used i.e. 8bp. No maximum limit has been set to include long repeats of functional importance. In case of imperfect repeats, minimum alignment score criteria are calculated by multiplying match weight of 2 with a minimum length of the locus for each repeat motif length ranging from 1-100.

### Interspersed Repeats: Transposable elements

Transposable elements constitute the major portions of the genomes. Expert curetted libraries consisting of several families and subfamilies of retrotransposons, DNA transposons are already available in databases like Repbase. *Oryza sativa* transposon sequence library prepared by REPEATMASKER software (http://www.repeatmasker.org/) in Repbase database has been used to search these curetted families in other rice genomes using blast with 10^-6^ e-value cut-off and 95% sequence identity and query coverage [47]. Parameters used in the present work for extraction of different classes of repeats have been tabulated in Table 2.

### Annotation and locations of the repeats

Annotations of repeats are parsed with respect to their locations in the genomes. For repeats occurring in the exons, UTRs and CDS regions, gene symbol, locus tag and product information of the gene have been used as the annotation of the repeat. Rice Annotation Project (RAP) [48] has taken the initiative to detect loci by mapping the sequences of transcripts and proteins to other monocot species and also to integrate annotation data from various resources like published literature, specialized databases consisting of annotations for several plant genomic elements e.g. microRNAs, small RNAs etc. As repeats are found to occur in the small and long RNA encoding genes, annotations from these databases can be used for the repeats.

### Statistically significant repeats

To find the statistical significance the extracted repeats of motif length 1-100 bp, comparisons have been done chromosome-wise across rice species by generating random genomes for each chromosome of every rice species. 1-sample t-test at significance level 0.05 has been conducted to check whether repeats are randomly distributed in the genomes or not.

### Comparisons among Oryza, Arabidopsis, and Brachypodium

Model dicotyledonous species *Arabidopsis* and another monocotyledonous wild grass *Brachypodium* have been selected for comparison with *Oryza*. Genomic parameters and repeat density parameters for both tandem and interspersed repeat classes have compared among all *Oryza* species with *Arabidopsis* and *Brachypodium.* To calculate whole genome size, individual sizes of all chromosomes have been added. The arithmetic mean of the %GC contents of all chromosomes is chosen for the %GC content comparison. The weighted mean of chromosome-wise gene densities per kbp has been computed. In case of calculating the %genic, %exonic and %coding region for the whole genome, respective amounts from each chromosome are added and then normalized by total genome size. While calculating repeat density parameters, repeat density in length parameter has been found same way as did for the %coding region calculation and for repeat density in number parameter weighted mean is used like gene density per kbp calculation.

### Searching repeats in the stress-related genes and association mining

As mentioned before, several experimental and computational studies have been cited in literature for identifying stress-related genes in rice and other crop species [49–52]. Many microarray-based transcriptomic studies also have been conducted and available in literature [25, 53–54] for stress associated gene detection. However, the available list is still not comprehensive and with rapid changes in global climate, more genes are affected and to be associated with stress tolerance mechanisms in rice. Nevertheless, positive (stress associated) and negative (uniformly expressed in any condition) sets have been constructed from available lists of genes reported in the published articles (Figure 2). Both positive and negative sets of genes are available for only *Oryza sativa* species. Hence, comparisons of repeat distributions have been performed for this species only. A nPMI based method [55] has been utilized for finding an association between stress and the repeats.

**Figure 2:**
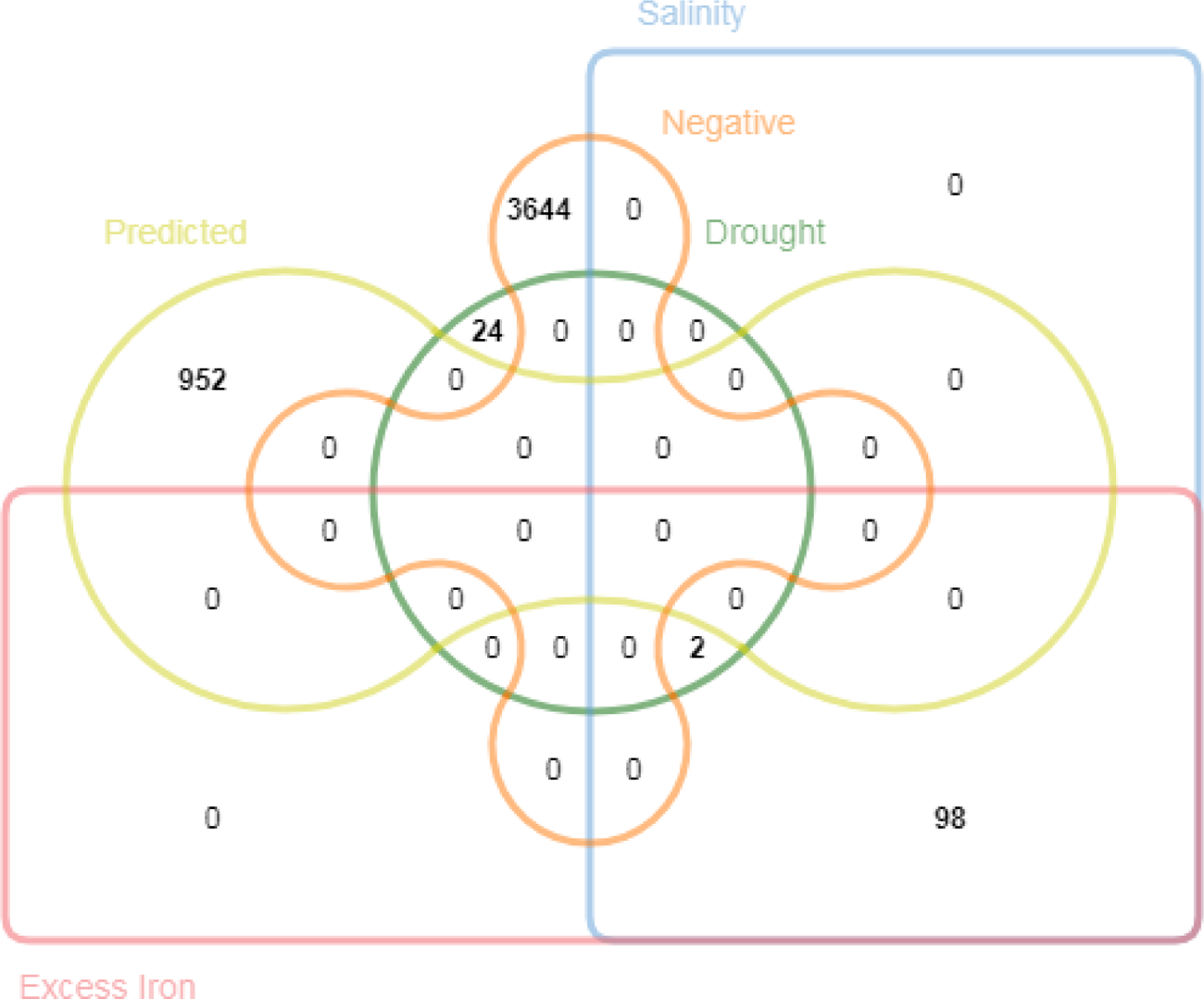
List of positive (stress related) and negative (housekeeping i.e. uniformly expressed under any condition). Venn diagram is created using jVenn web tool [56]. Red: Excess iron; Cyan: Salinity; Green: Drought; Yellow: Predicted stress genes; Orange: Negative or Housekeeping; Least common gene list is observed here.

### Building positive set

A positive set of stress associated genes in rice has been collected from several published articles. Recently, Lakra et al. have reported 190 salinity stress-related proteins which are identified from proteomics study of contrasting rice genotypes [49]. In another study by Finatto et al., total 2525 rice genomic loci have been reported to be influenced by excess iron stress [25]. A well-known database that contains curetted drought stress-related genes for a number of plant species including *Arabidopsis,* rice etc. has been developed by Alter et al which contains 30 drought stress-related genes for rice [50]. Finally, 2692 genomic loci have been selected as gold standard (Set A) stress gene set. Many exclusive theoretical studies have been performed to identify plant stress-related genes which include predictions using orthology by Prabha et al. [52], transcription factor interaction network [51, 54], gene expression analysis [57] etc. Total 952 genomic loci have been collected from aforementioned theoretical studies to construct predicted stress-related genes (Set B) for further analysis.

### Constructing negative set for stress (NSS)

To date, it is still difficult to identify housekeeping genes (HK) those are uniformly expressed under any conditions. Very few rice genes have been reported as housekeeping genes. But the number is very low for comparative analysis of repeat distributions. To overcome this inadequacy, predicted housekeeping genes from Rice Expression Database (RED) [58] have been used for comparative analysis. RED is a comprehensive database that stores gene expression profiles from 284 high-quality RNA-Seq experiments performed under various treatments and a wide range of growth stages. The database enlists a set of housekeeping genes that are uniformly expressed under any condition. On the basis of statistical parameter ‘τ’ [59] which indicates the extent of expression breadth, HK genes have been sorted in ascending order. To balance the positive and negative classes for unbiased prediction equal no. (As of positive gene) of top HK genes having low ‘tau’ value and does not overlap with the positive set, have selected as the negative set. List of the positive and negative gene lists have been given in the Supplementary Table file.

## Results & discussions

### Rice genomes and availability of annotations

As mentioned earlier, rice is a diploid species which has 12 pairs of chromosomes with size much larger than *Salmonella* chromosome (approximately 80 times). It is the first crop genome that is sequenced after successful sequencing of the flowering model plant *Arabidopsis* which is much smaller in size than *Oryza* (approximately one-third of rice genome). Initiatives have been taken from several institutes for sequencing and assembling the aforementioned rice species. Among them, Arizona Genomics Institute (AGI) have sequenced and assembled 5 *Oryza species* including *O. barthii*, *O. glaberrima*, *O. glumaepatula*, *O. nivara* and *O. punctata*. Sequencing and assembling of *O. brachyantha*, *O. indica,* and *O. rufipogon* have been done in the three different institutes of China namely Institute of Genetics and Developmental Biology (IGDB), Chinese Academy of Sciences (CAS), Beijing, Beijing Genome Institute (BGI) and Shanghai Institutes for Biological Sciences, CAS respectively. Model rice species *O. sativa* has been sequenced and assembled collaboratively by *MISSISSIPPI STATE* University (MSU) and International Rice Genome Sequencing Project (IRGSP) members [60]. In the year 2000, India becomes a part of IRGSP and took the challenge of sequencing chromosome 11 jointly ventured by University of Delhi South Campus (UDSC), the National Research Centre on Plant Biotechnology (NRCPB) and the Indian Agricultural Research Institute (IARI), New Delhi. Recently, Indian scientists have developed markers and detected fragrance alleles in the rice varieties [61–62].

Two major initiatives have been taken by MSU and Rice Annotation Project (RAP) for annotating the model species *Oryza sativa* after completion of its sequencing in 2004 [63] using various *ab initio* methods and EST/mRNA alignments to produce the final gene models. MSU gene models have been also generated using automatic annotation pipeline which includes several tools like FGENESH, Genemark, Genscan, GeneSplicer, tRNAScan etc. Complete annotation and cross-references between two gene models are available at RAP database (http://rapdb.dna.affrc.go.jp) which is adopted by several databases like NCBI, Gramene etc. Annotations for other rice species have been performed by the corresponding sequencing groups either individually or in collaboration with other groups. In summary, protein-coding gene annotations have been done by evidence-based MAKER-P genome annotation pipeline mainly. Finding annotations for non-coding RNA genes have been executed using tRNAScan program [64] and transposable elements have been detected using RepeatMasker tool. For the majority of the rice species, genome assemblies and annotations are not finalized yet which is reflected from the presence of contigs in the sequences and very few number of RNA associated genes in the annotations compare to *O. sativa* and *O. indica* (Table 3).

**Table 3:** Availability of protein-coding, RNA associated genes and transcripts from Ensembl Plants release 37 (http://plants.ensembl.org/)

| No. | Species | Protein Coding Genes | RNA associated genes | Transcripts |
| --- | --- | --- | --- | --- |
| 1 | *Oryza barthii* | 34575 | 3212 | 44891 |
| 2 | *Oryza brachyantha* | 32038 | 2560 | 34598 |
| 3 | *Oryza glaberrima* | 33164 | 41415 | 74579 |
| 4 | *Oryza glumaepatula* | 35735 | 3967 | 50860 |
| 5 | *Oryza sativa indica* | 40745 | 48978 | 89723 |
| 6 | *Oryza nivara* | 36313 | 2428 | 50788 |
| 7 | *Oryza punctata* | 31762 | 6029 | 47089 |
| 8 | *Oryza rufipogon* | 37071 | 3412 | 51004 |
| 9 | *Oryza sativa japonica* | 35679 | 56313 | 98663 |

### Comparison of genomic parameters in Oryza species

Model species *Oryza sativa* has the genome of size 373.25 Mbp and average %GC content of 43.6. As shown in Figure 3A-3B, variations in genome sizes and % GC content indicate its ability to adapt to diverse environmental conditions [65]. In spite of low variance in gene density per kbp across 9 *Oryza* species (variance 0.21) (Figure 5.3C) percentages of genic regions are highly varying (Figure 3D). %Exonic and %CDS regions are also varying across 9 rice species (Figure 3E-3F). Model species *O. sativa* has the 0.244 genes per kbp of the genome which is higher than all other species except *O. glaberrima.* Another important observation is that all the domesticated rice species i.e. *O. glaberrima*, *O.indica*, *O. punctatata* and *O. sativa* have lower percentages of the genic region than their wild wild progenitors which include African wild species *O. barthii*, *O. brachyantha*, South American wild species *O. glumaepatula* and Asian wild rice *O. nivara* and *O. rufipogon*. This might be due to the fact of reduction of transposable elements in genic regions across the domesticated species [66]. However, all of the species possess comparable percentages of exonic and protein coding regions.

**Figure 3:**
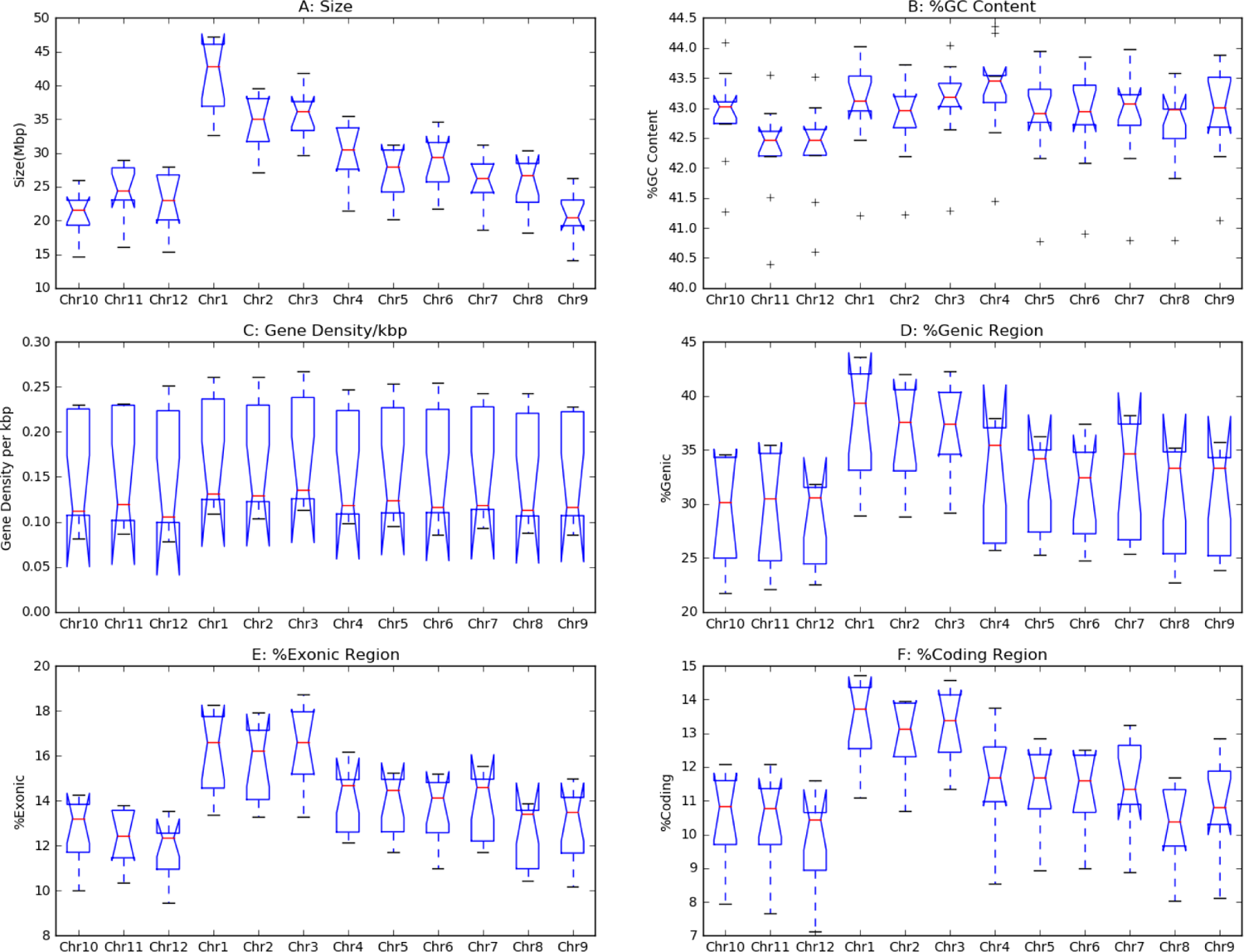
Comparison of genomic properties across *Oryza* species; A: Size in Mbp; B: %GC content; C: %Genic Region; D: Gene density per kbp; E: %Exonic Region; F: %Coding Region. Along X-axes of each subplot, chromosomes with number have been shown. For example, Chr1 abbreviates for chromosome number 1.

### Different classes of repeats and their distributions in chromosomes

Both perfect and imperfect tandem repeats are abundant in rice chromosomes covering more than 5% of the genome which include both microsatellites and minisatellites present inside genes, intergenic regions and transposable elements. Model species *O. sativa* possess maximum amount of repeats (*D_L_* ∼ 6.4%) compare to other rice species followed by *O. rufipogon* (*D_L_* ∼ 5.9%) and *O. indica* (*D_L_* ∼ 5.7%). Similarly, in the same species, *O. sativa* maximum numbers of repeats are found to occur per Mbp (*D_N_* ∼ 4182 repeats/Mbp) followed by *O. rufipogon* (*D_N_* ∼ 4060 repeats/Mbp), *O. barthii* (*D_N_* ∼ 3936 repeats/Mbp) and *O. indica*(*D_N_* ∼ 3929 repeats/Mbp) (Figure 4-5). On average, 3906 tandem repeats are occurring per Mbp of the rice genome. In case of interspersed repeats, only 1.5% of the genome is covered with this kind of repeats (median *D_L_* ∼ 1.51%) and their number of occurrences is much lower than that of tandem repeats (median *D_N_* ∼ 30 repeats/Mbp). Assembling and finishing off the rice genomes have not fully completed yet (95% coverage) and contains gaps and uncharacterized sequences. Hence, there remains a possibility that these numbers might change after gap filling and inclusion of uncharacterized parts into rice chromosomes as seen in the recently assembled rice genome with 99% coverage [40].

**Figure 4:**
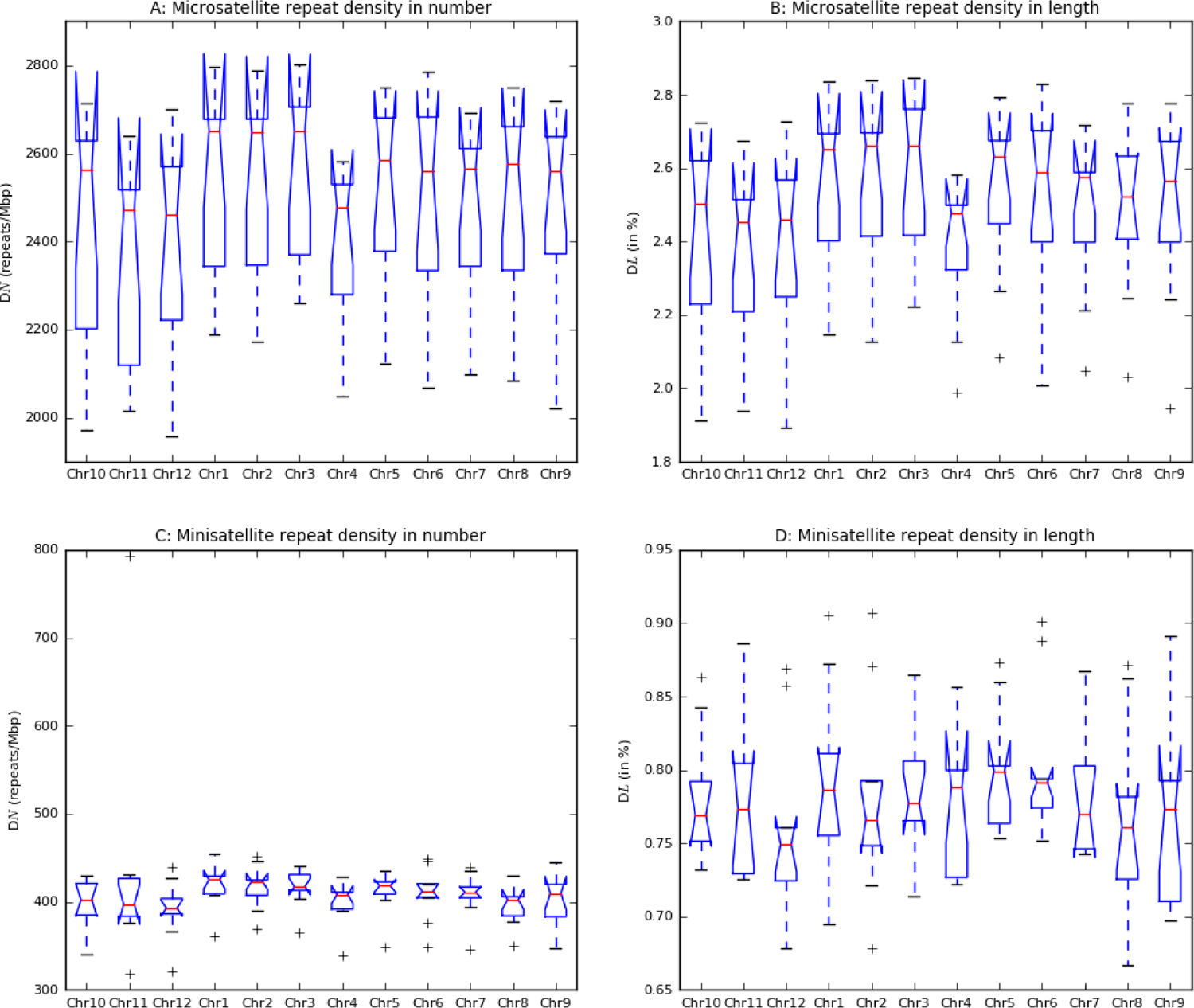
Comparing distributions of different classes of perfect repeats in 9 *Oryza* genomes. A: Microsatellite repeat density in number (*D_N_*); B: Microsatellite repeat density in length parameter (*D_L_*); C: Minisatellite repeat density in number (*D_N_* parameter) and D: Minisatellite repeat density in length parameter (*D_L_*).

**Figure 5:**
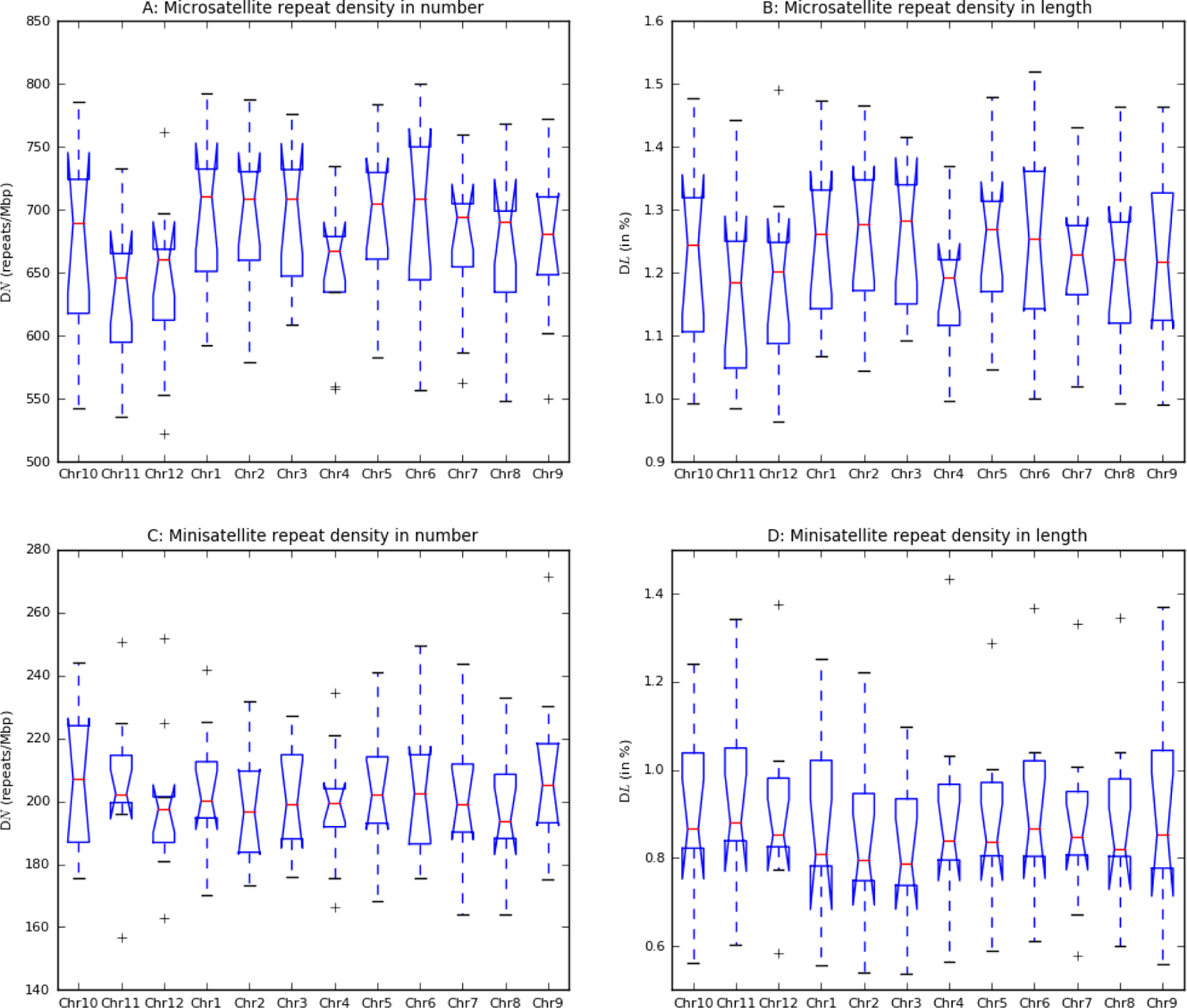
Comparing distributions of different classes of imperfect repeats in 9 *Oryza* genomes. A: Microsatellite repeat density in number (*D_N_*); B: Microsatellite repeat density in length parameter (*D_L_*); C: Minisatellite repeat density in number (*D_N_* parameter) and D: Minisatellite repeat density in length parameter (*D_L_*).

### Microsatellites

Microsatellites are the most predominant class of repeats in rice. On average more than 2500 perfect microsatellites are present per Mbp of rice genome. *O. sativa* has the highest *D_N_* for perfect repeats (*D_N_* ∼ 2728 repeats/Mbp) and *O. punctata* has the lowest number (*D_N_* ∼ 2097 repeats/Mbp). Among the 9 species, chromosome 1 has the highest *D_N_* value (median 2652 repeats/Mbp) whereas chromosome 12 has the least (median 2461 repeats/Mbp) (Figure 4A). In terms of *D_L_* parameter, approximately 2.5% of genome is covered by perfect microsatellites with very low variation across species (< 0.05). Their abundance is also equal to 2.5% of a chromosome (variance of 0.006 across chromosomes) (Figure 4B). *O. sativa* followed by *O. rufipogon* have more than 2.7% of their genome covered by perfect microsatellites whereas *O. punctata* has the least value of coverage i.e. 2.04%. Imperfect microsatellites have occurred less frequently than perfect ones (*D_N_* ∼ 690 repeats/Mbp) with large variations across species (Figure 5A). Approximately 1.2% of the rice genome is covered by imperfect microsatellites with a variance lower than 0.02. The coverage of imperfect microsatellites in each chromosome is 1.2% too and variance is below 0.001 (Figure 5B). So, microsatellites are almost equally distributed in the chromosomes but highly varying in numbers and might be of functional importance.

### Minisatellites

Minisatellites are almost one fifth (in terms of numbers) of the abundance of the microsatellites. Perfect minisatellites are frequent (median *D_N_* ∼ 413 repeats/Mbp) compare to imperfect ones (median *D_N_* < 200 repeats/Mbp) (Figure 4C-5C) but covers little less portion of the genome than polymorphic minisatellites (mean *D_L_* ∼ 0.8% and 0.9% respectively) (Figure 4D-5D). *O. sativa* has the maximum numbers of minisatellites repeats per Mbp whereas *O. rufipogon* has more minisatellites coverage in the genome. Like perfect microsatellites, perfect minisatellites are also equally distributed across chromosomes (mean *D_L_* ∼ 0.7% per chromosome and variance ∼ 0.0001) and not varying too much across species (variance < 0.05). Similar observations have been found in case of imperfect minisatellites also. In terms of *D_N_* parameter, perfect minisatellites are little less varying across species compare imperfect ones. So, tandem repeats are equally distributed across chromosomes but like microsatellites minisatellites are also varying in number per Mbp of the genome.

### Interspersed repeats

As mentioned earlier in the materials and method section, interspersed repeats are collected from Repbase which is the reference database used by the tools for identifying interspersed repeats in the eukaryotic genomes. This particular database is chosen because it is a curetted, regularly updated resource and has utilized a specialized scheme for classifying and annotating transposable elements which constitutes the major part of the interspersed repeats in any eukaryotic species. In the database, transposable elements are listed for the model species *Oryza sativa* only (Figure 6). Using programs like Blast, these reference sequences can be mapped to the other rice species for finding highly homologous transposable elements that can perform similar functions across rice species. Transposable elements can occur in multiple copies in a genome. Hence, ‘-max_target_seqs’ and‘-max_hsps’ parameters of blastn program have been set as default i.e. 500 and all HSPs. But to find highly similar sequences with highly similar functions, more than 95% sequence identity and query coverage with an expected value of 10^-6^ filters have been applied. Though, it have filtered and reported much fewer amounts of transposable elements (maximum *D_L_* of 7.75% in *O. sativa*) than earlier reports (∼37.5%) but the extracted elements are highly reliable for stress association analysis and prediction (Figure 7).

**Figure 6:**
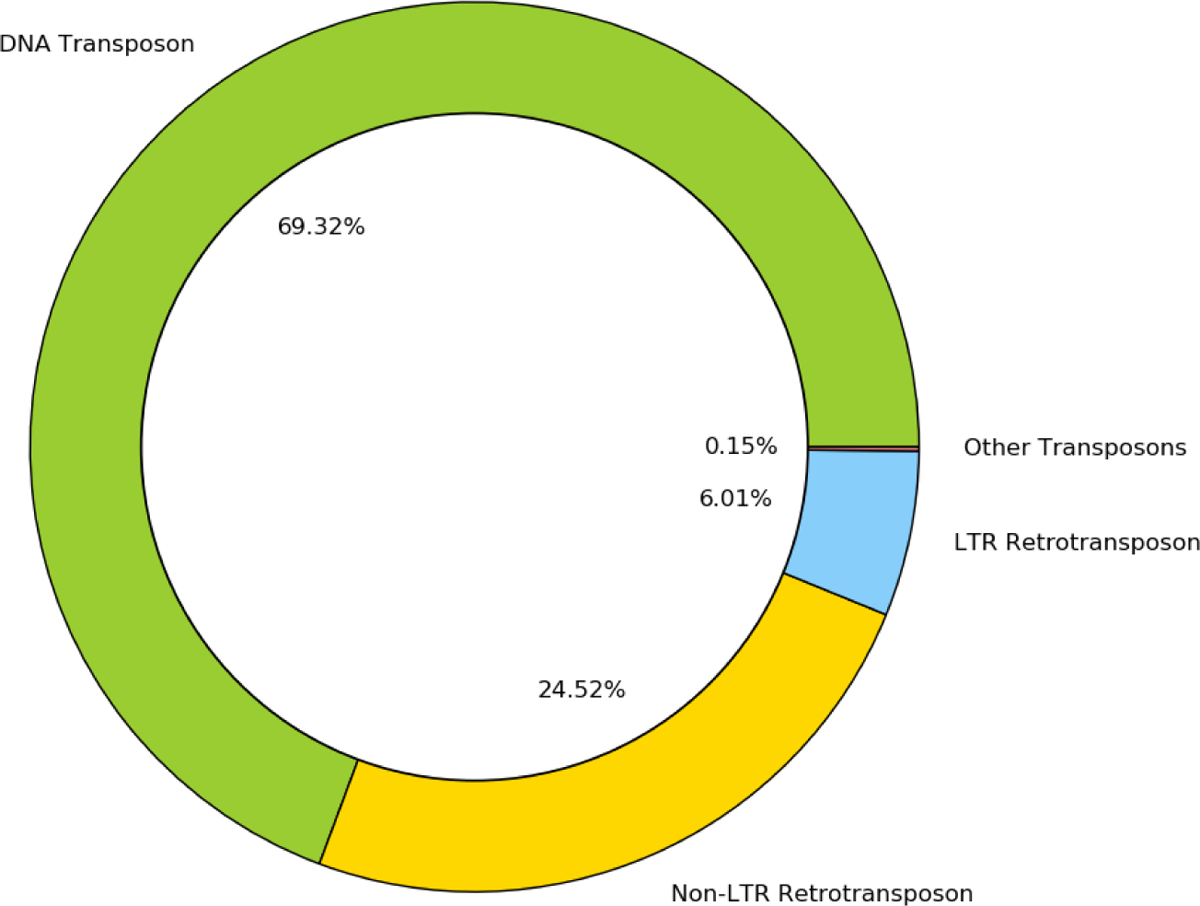
Different classes of Interspersed Repeats in *Oryza sativa* in Repbase v20.05.Green: DNA Transposons; Blue: LTR Retrotransposons; Yellow: Non-LTR Retrotransposons; Black: Other transposons.

**Figure 7:**
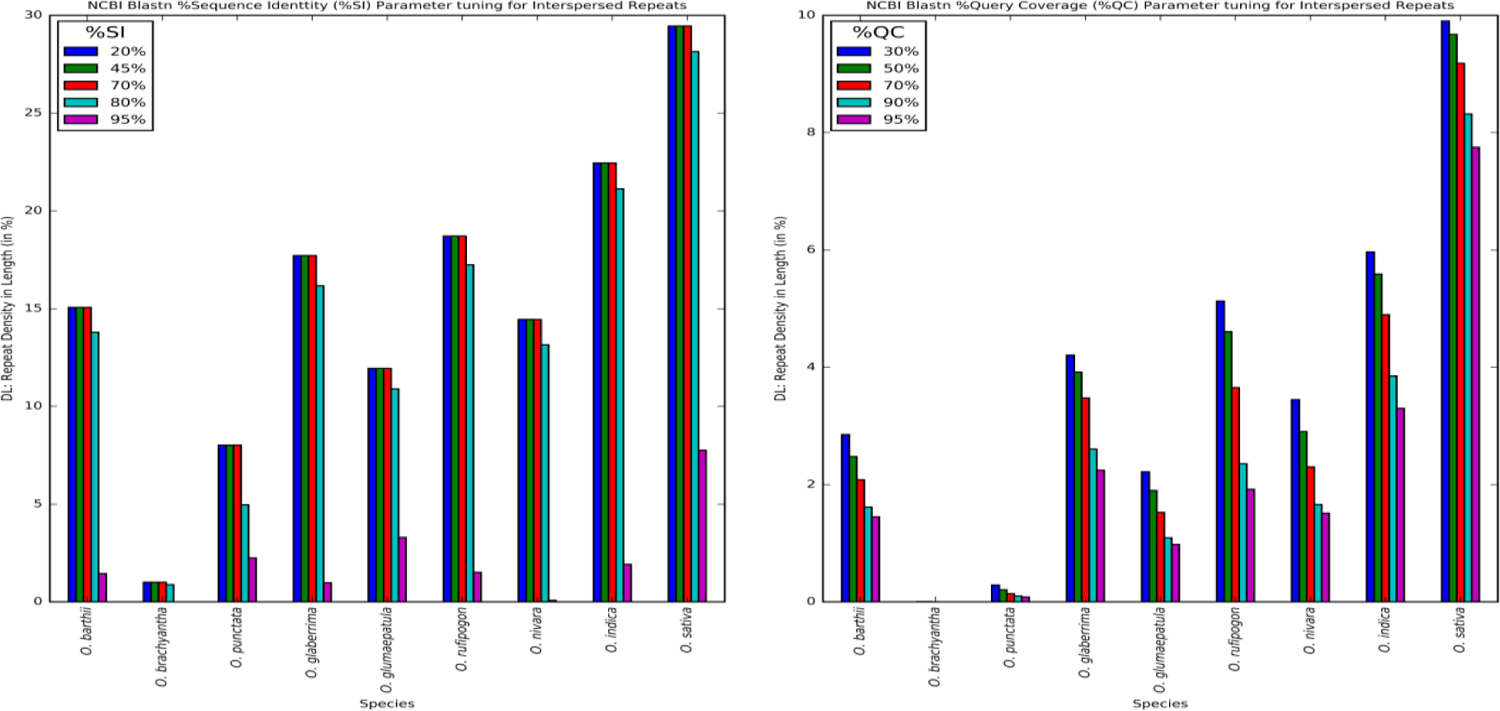
Comparison of blast parameters for interspersed repeat extraction from rice species. A: Percent sequence identity and B: Percent query coverage

As expected, *O. sativa* possess maximum number of transposable elements (*D_N_* ∼ 75 repeats/Mbp) and *O. brachyantha* has less than 1 repeat/Mbp (Figure 8A-8B). Effect of extreme strict criteria for the selection of highly reliable interspersed repetitive elements is reflected in the current result.

**Figure 8:**
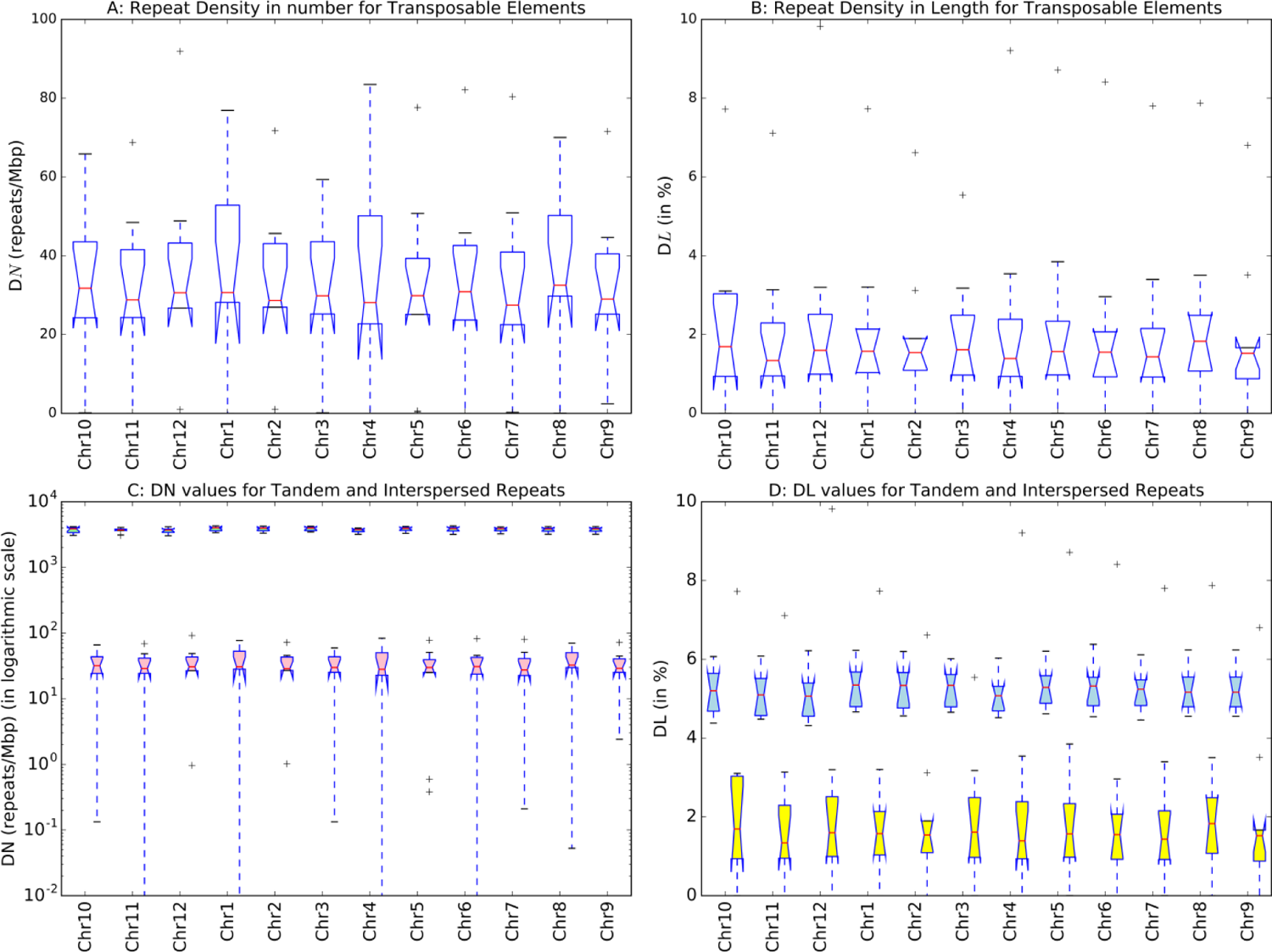
Distribution of Interspersed Repeats in Oryza sativa and comparison of Tandem and Interspersed Repeats in Oryza sativa on the basis of repeat density in number parameter. Pink and yellow are interspersed and green and blue are for tandem repeats

### Statistically significant repeats

In the frequency analysis of tandem repeats of motif lengths ranging from 1-100 bp including both perfect and imperfect repeats, the very first observation is that the perfect tri-meric repeats are occurring significantly (1sample t-test *p*-value < 0.05) more frequently than the other motifs followed by monomeric and di-meric repeats. Most striking observation is that perfect minisatellites do occur in the real genomes but absent in the random genomes as expected (Figure 9-10 and supplementary Figure 1A and Figure 1B). All repeats of motif lengths 1-6 bp are occurring significantly the rice genomes (1sample t-test *p*-value < 0.05). Imperfect repeats of motif lengths 1 to 100 nucleotides are present in the rice genomes but with less number per Mbp (Figure 11-12) than perfect ones and also statistically significant (1sample t-test *p*-value < 0.05). Unlike perfect minisatellites, imperfect minisatellites have been occurred significantly different from their occurrence in the random genomes. Both perfect and imperfect minisatellites are less frequent than the occurrence of microsatellites.

**Figure 9:**
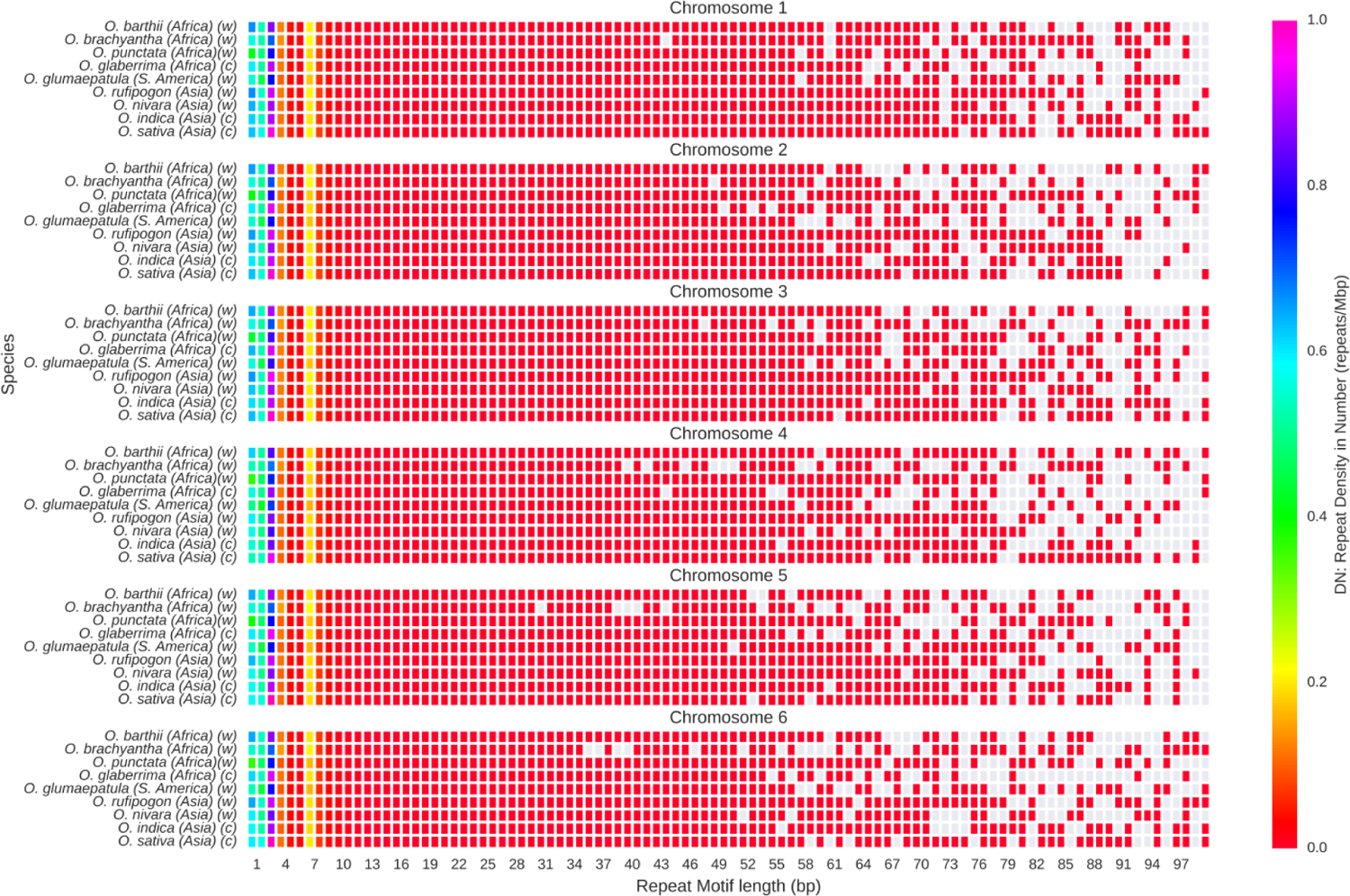
Distribution of perfect repeats with different motif lengths (1-100) across *Oryza* genomes. Chromosomes 1 to 6 have been shown here. Color bar represents repeat density in number (*D_N_* parameter in repeats/Mbp) parameter. From red to magenta denotes lower values to higher values.

**Figure 5:**
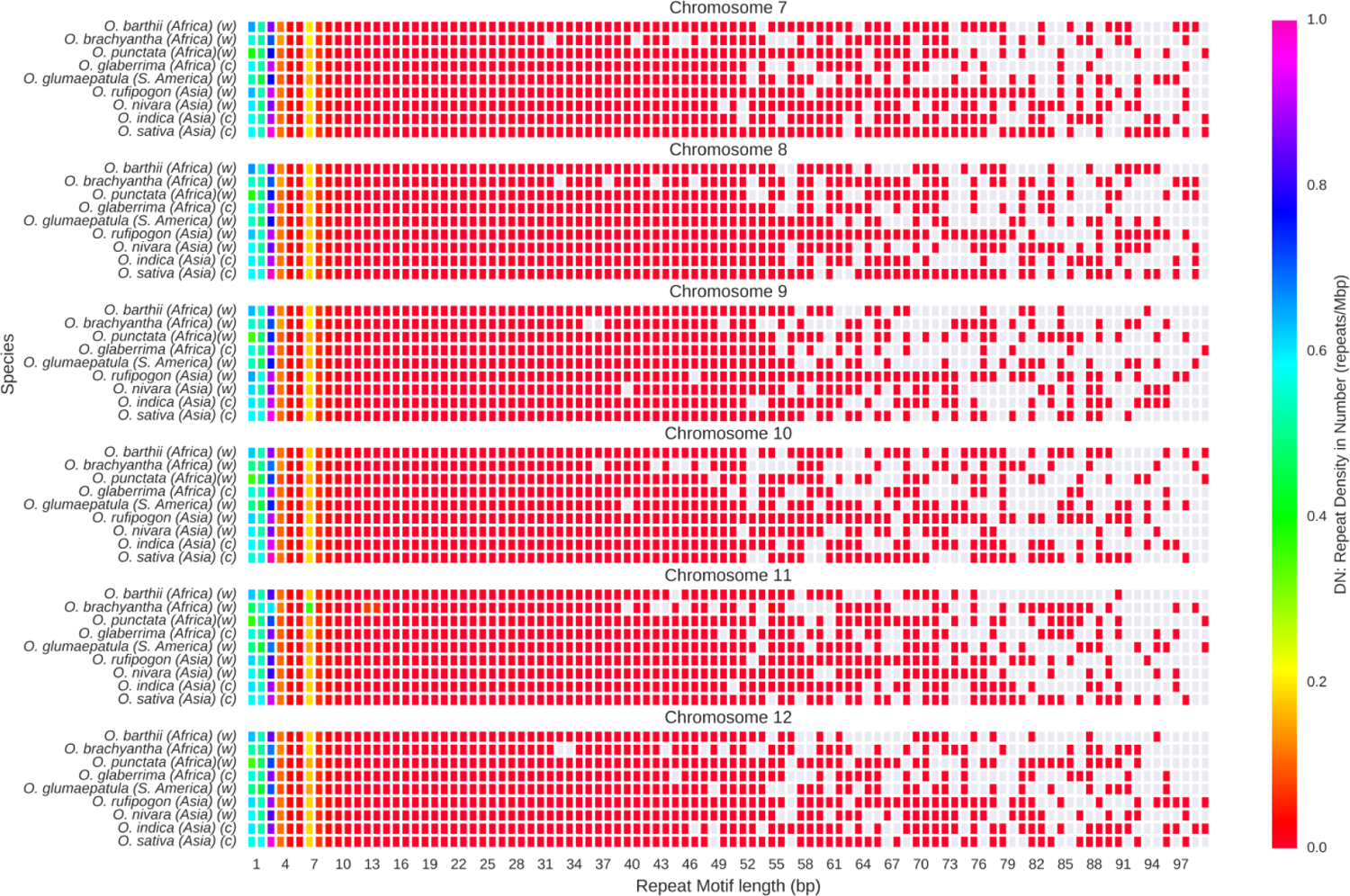
Distribution of perfect repeats with different motif lengths (1-100) across Oryza genomes. Chromosomes 7 to 12 have been shown here. Color bar represents repeat density in number (DN parameter in repeats/Mbp) parameter. From red to magenta denotes lower values to higher values.

**Figure 11:**
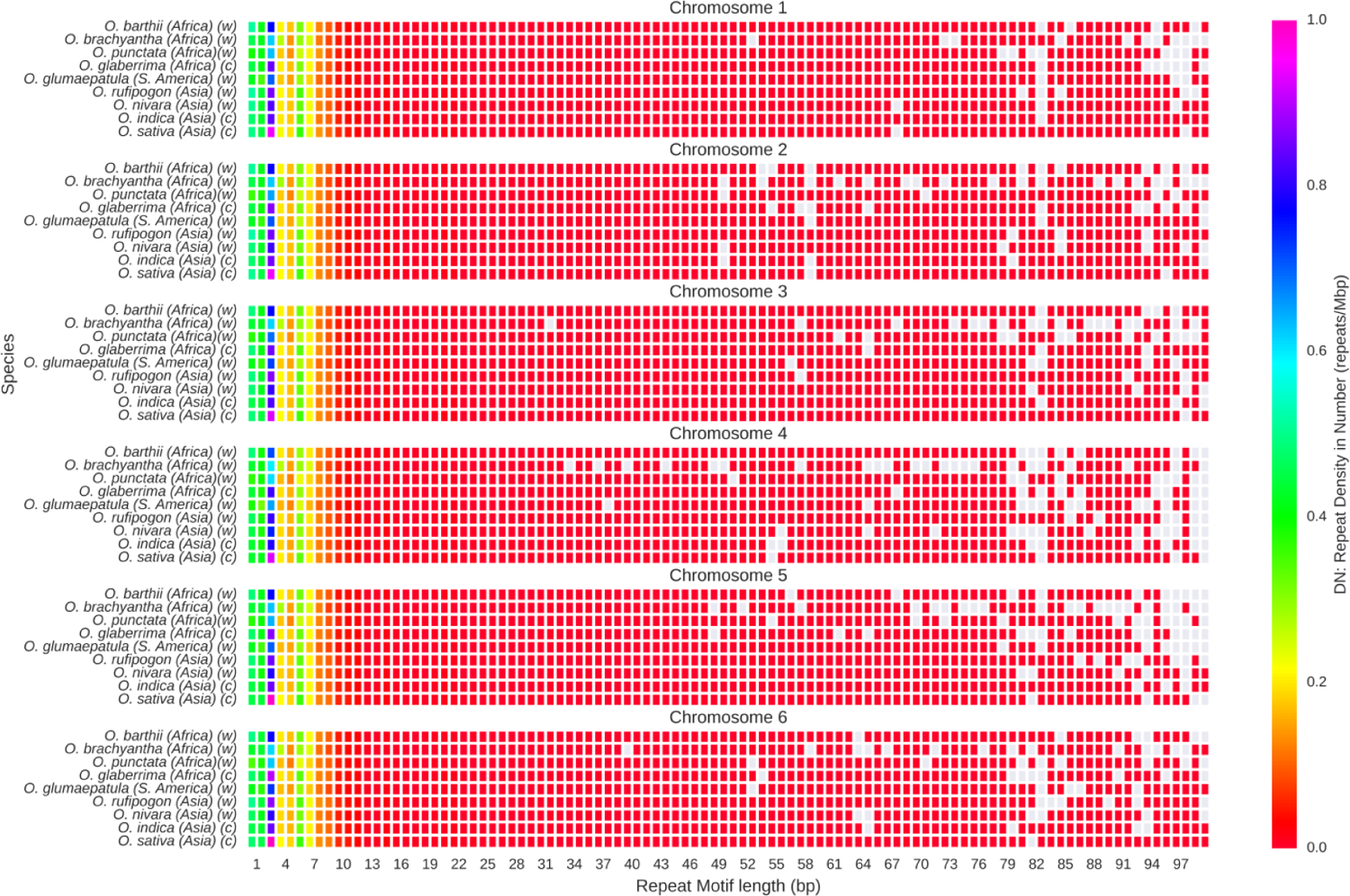
Distribution of imperfect repeats with different motif lengths (1-100) across *Oryza* genomes. Chromosomes 1 to 6 has been shown here. Color bar represents repeat density in number (*D_N_* parameter in repeats/Mbp) parameter. From red to magenta denotes lower values to higher values.

**Figure 12:**
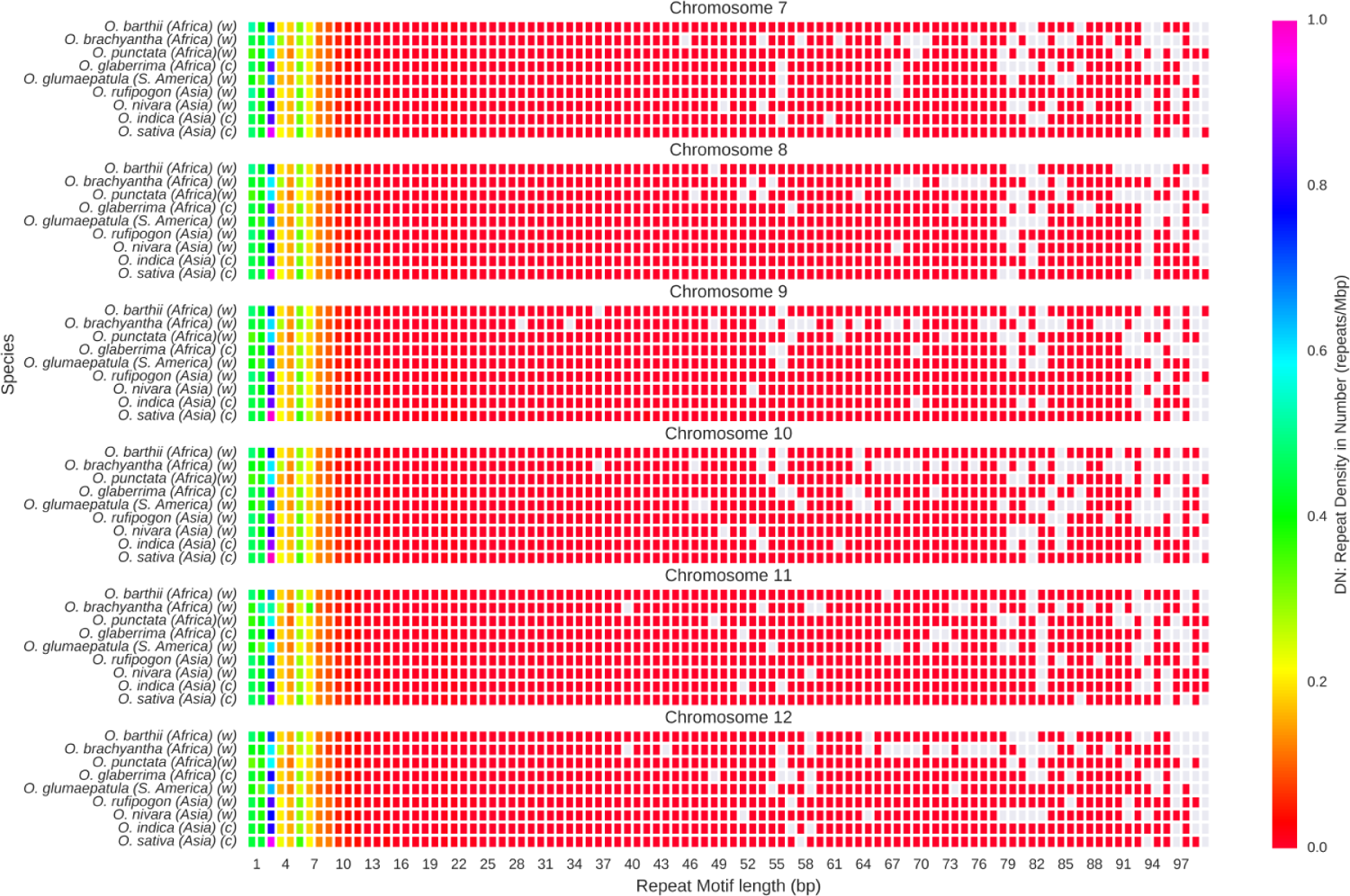
Distribution of imperfect repeats with different motif lengths (1-100) across *Oryza* genomes. Chromosome 7 to 12 has been shown here. Color bar represents repeat density in number (*D_N_* parameter in repeats/Mbp) parameter. From red to magenta denotes lower values to higher values.

Heptameric repeats are more ubiquitous than the other minisatellites ranging from length 8 bp to 100 bp. These observations signify that trimeric repeats have ubiquitously occurred in the genomes compared to other microsatellites [67]. A major portion of the rice genome is not the coding region. So, the presence of trimeric repeats in excess is not only due to their relation with codons but also they must have some definite functional importance. Several earlier reports have shown that both micro- and minisatellites regions are prone to methylation leading to epigenetic regulations of genes and genomes and their crucial roles in various developmental processes such as the development of seeds, embryos and gametophytes, controlling flowering time and stress responses [68].

### Occurrence of repeats in different genomic locations

As shown in series of Figures 13-18, perfect and imperfect tandem repeats are ubiquitous in different genomic locations of rice genomes including genic, intergenic, exons, introns, coding and untranslated regions.

**Figure 13:**
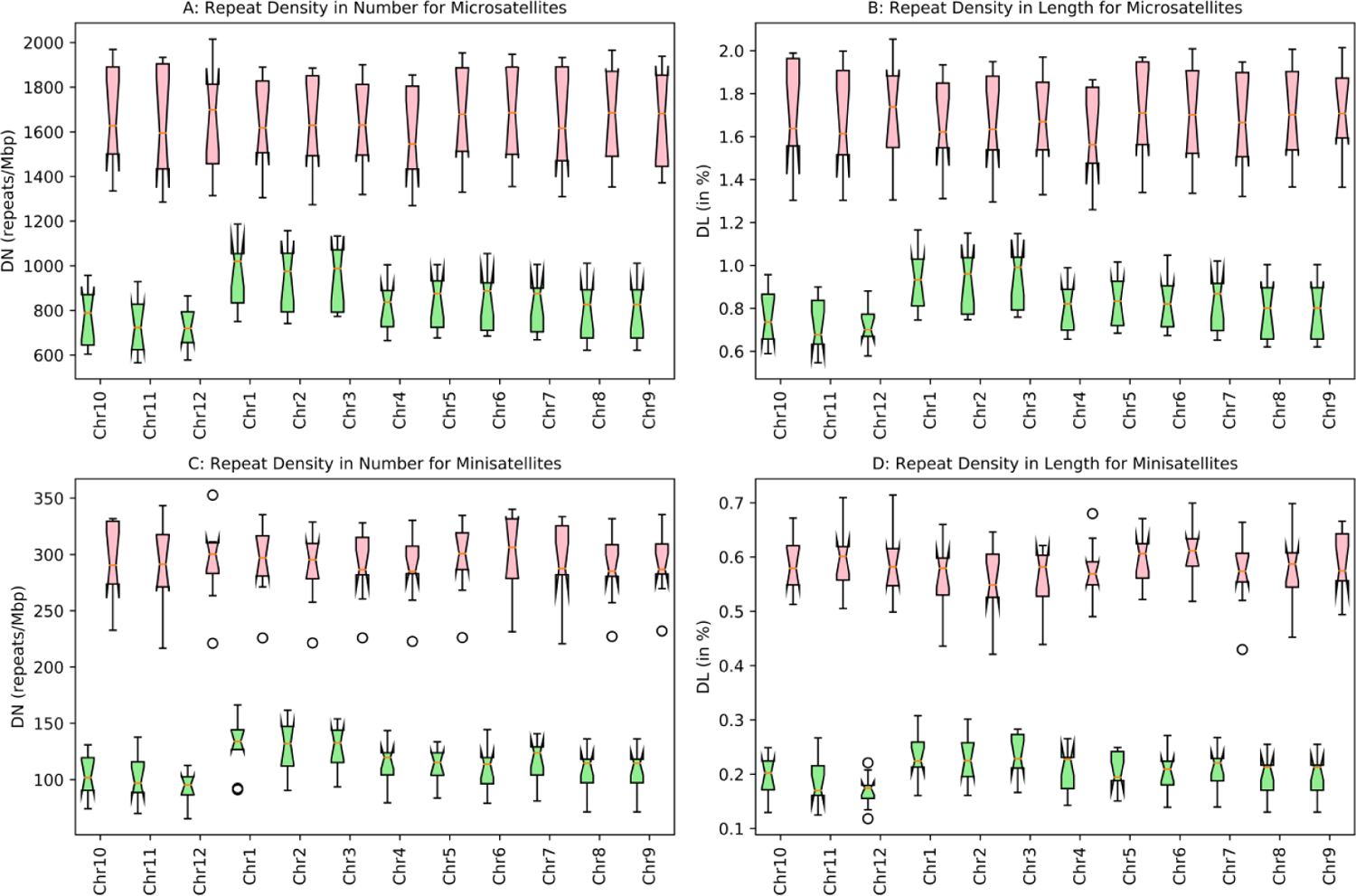
Chromosome-wise distribution of Perfect Tandem Repeats (Microsatellites and Minisatellites) in Genic (Green) and Intergenic regions (pink)

Repeats in genic and intergenic regions

On average, 884 perfect microsatellite repeats/Mbp are occurring in the genic regions which are almost half than that of intergenic regions (median value 1640 repeats) (Figure 13A). *O. rufipogon* has the maximum number of repeats (*D_N_* ∼ 1037 repeats/Mbp) followed by *O. nirvara* (*D_N_* ∼ 1018). *O. sativa* has the *D_N_* value of 884 repeats/Mbp but contains the maximum number of perfect microsatellites in the intergenic regions (*D_N_* ∼ 1909 repeats/Mbp).

Perfect microsatellites in the genic and intergenic regions have covered approximately 0.8% and 1.6% of the genome which are also uniformly distributed across the chromosomes (variance < 0.006) (Figure 13A-13B). Perfect minisatellites are also frequent in the intergenic regions (median *D_N_* ∼ 296 repeats/Mbp) compare to genic regions (median *D_N_* ∼ 114 repeats/Mbp) and their numbers are varying across chromosomes too (Figure 13C-13D). In terms of repeat density in length parameter, genic and intergenic repeats cover approximately 0.2% and 0.6% of the genome respectively. Like microsatellites, perfect genic and intergenic minisatellites are also equally distributed in the chromosomes.

Unlike perfect microsatellites in the genic and intergenic regions, polymorphic microsatellites are much less frequent in numbers per Mbp (median value of ∼232 and ∼450 repeats/Mbp respectively) covering ∼0.4% and ∼0.9% of the genome respectively having nearly equal contributions from each chromosome (Figure 14A-14B). Polymorphic minisatellites are less abundant than microsatellites in genic and intergenic regions (median *D_N_* ∼ 53 and 144 repeats/Mbp respectively) covering 0.2% and 0.7% of the genomes respectively (Figure 14C-14D). Almost equal contributions from each chromosome have been observed in this case also.

**Figure 14:**
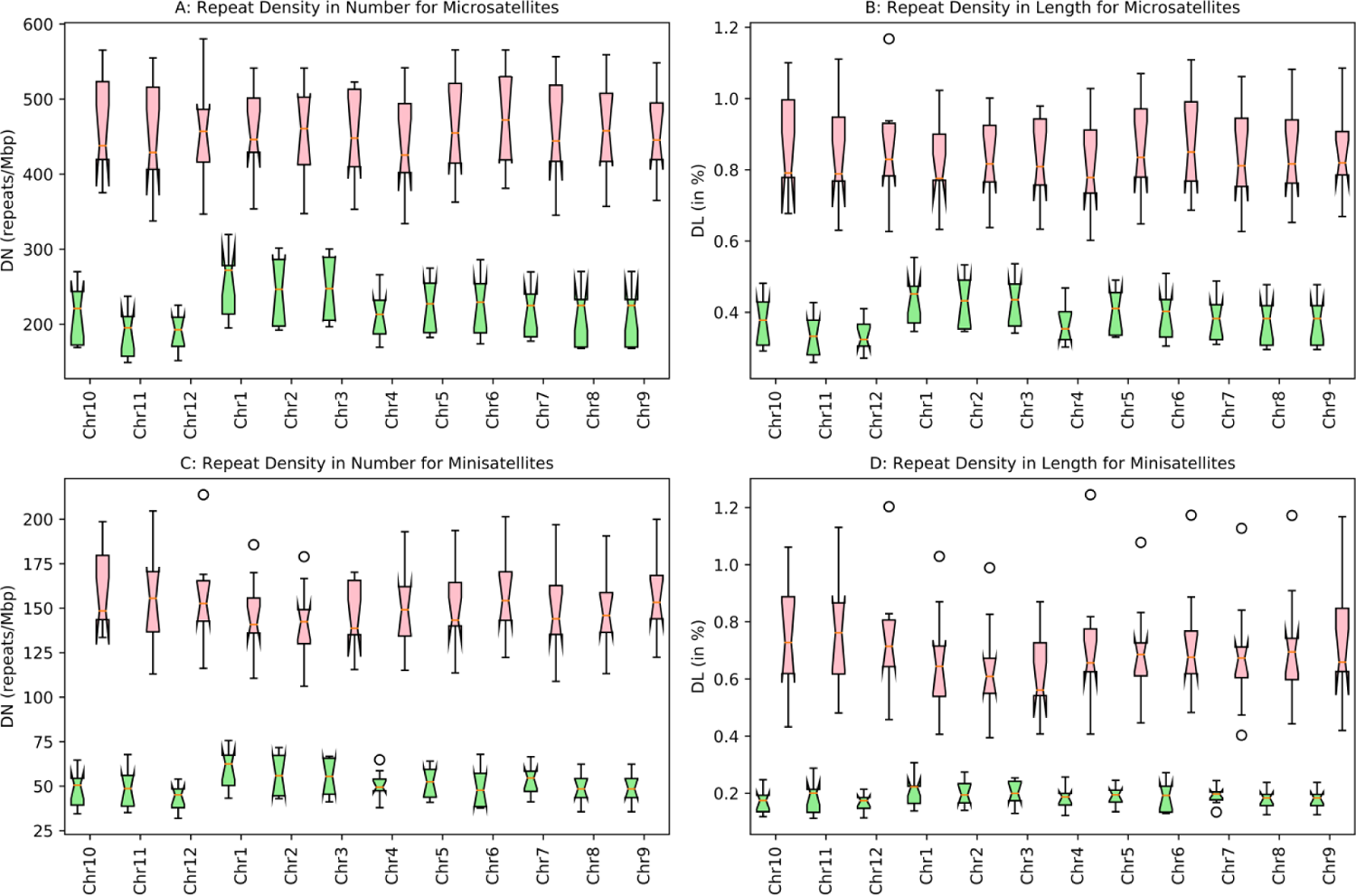
Chromosome-wise distribution of Imperfect Tandem Repeats (Microsatellites and Minisatellites) in Genic (Green) and Intergenic regions (pink)

### Repeats in exonic and intronic regions

Inside the genic regions, repeats are ubiquitously found both in the exonic and intronic parts of the genome. *D_N_* values for perfect microsatellites in exons and introns are comparable (median *D_N_* ∼ 479 and 422 repeats/Mbp respectively) (Figure 15A) and covering approximately 0.4% of the genomes in both cases (Figure 15B). In the case of imperfect microsatellites, their occurrences are slightly greater in the exonic regions (median *D_N_* ∼ 140 repeats/Mbp) than intronic regions (median *D_N_* ∼ 100 repeats/Mbp) (Figure 16A). A similar observation has been found in case of repeat density in length parameter also. Imperfect microsatellites in the exonic regions cover only 0.23% of the genome which is little higher than their presence in the intronic region (*D_L_* ∼0.16%) (Figure 16B). Perfect minisatellites are more abundant in intronic regions (*D_N_* ∼ 80 repeats/Mbp) compare to exons (45 repeats/Mbp) (Figure 15C). Exonic perfect minisatellites cover only 0.06% of the genome whereas intronic repeats cover approximately 0.14% of the genome i.e. more than double than that of perfect ones (Figure 15D). Percentages of occurrences of imperfect minisatellites in the exonic and intronic region are comparable (*D_L_* values ∼ 0.07% and 0.11% respectively) (Figure 16D). Similar finding has been observed in case of repeat density in number parameter too. DN values for imperfect minisatellites in exons and introns are 24 and 30 repeats/Mbp respectively. (Figure 16C). Surprisingly, *O. nivara* has maximum number of perfect and imperfect microsatellites than the other species in both regions. *O. sativa* has maximum number of repeats per Mbp in the exonic regions and *O. rufipogon* possess the maximum values in intronic regions.

**Figure 15:**
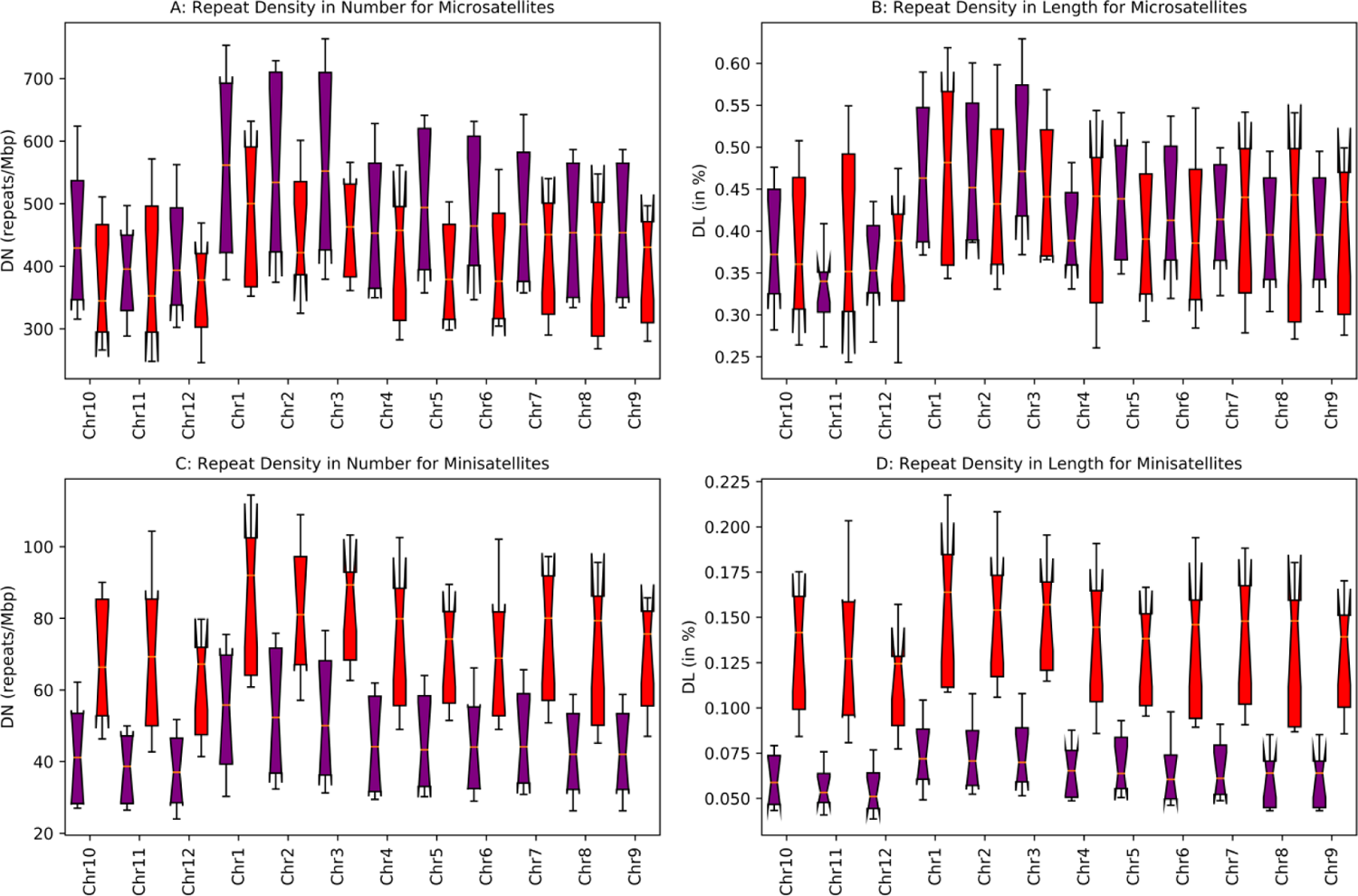
Chromosome-wise distribution of Perfect Tandem Repeats (Microsatellites and Minisatellites) in exons (Purple) and introns regions (Red)

**Figure 16:**
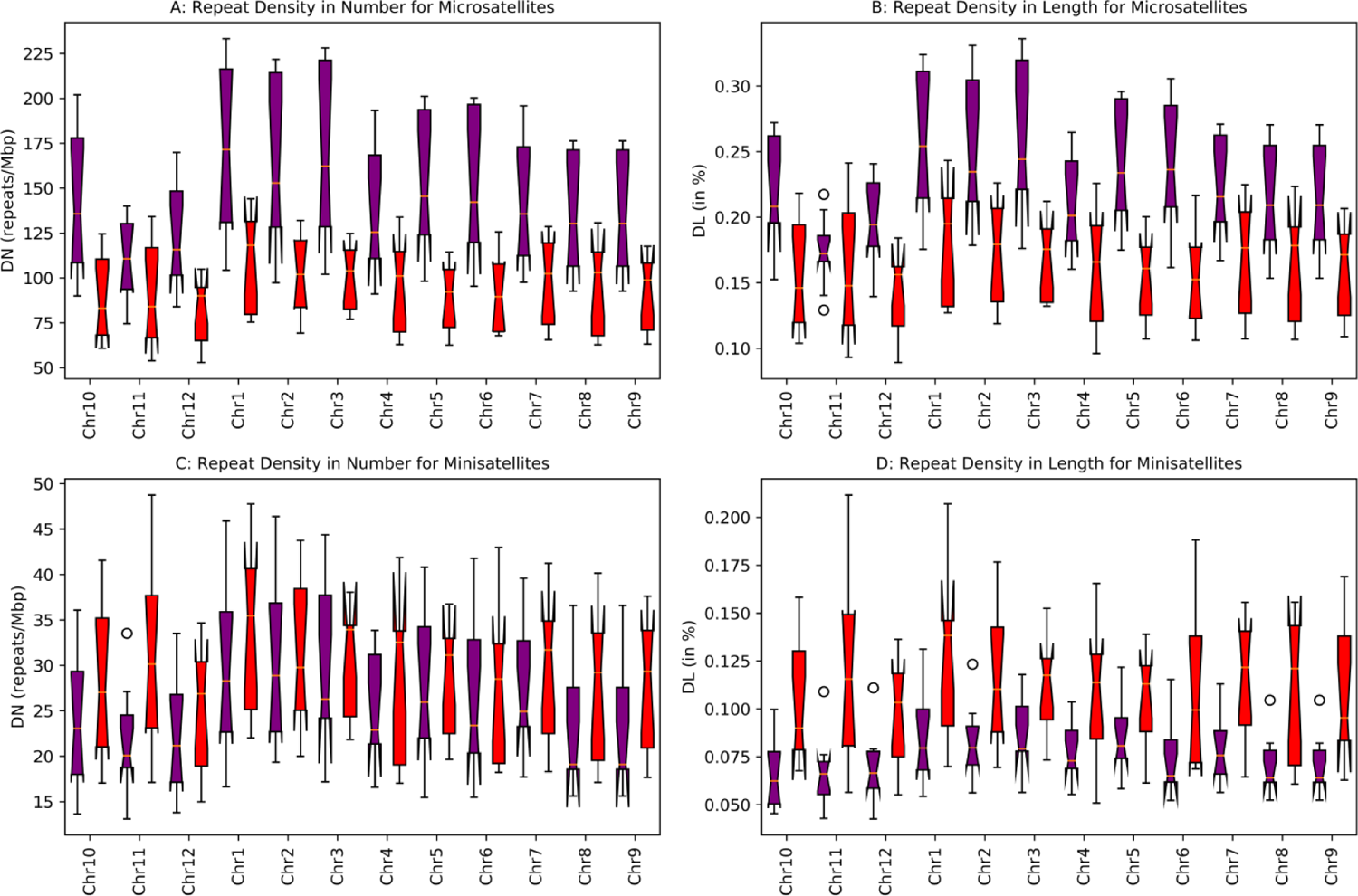
Chromosome-wise distribution of Imperfect Tandem Repeats (Microsatellites and Minisatellites) in exons (Purple) and introns regions (Red)

### Repeats in coding regions and untranslated regions

Perfect microsatellite repeats more occur in the coding regions than the untranslated regions. Repeat density in number parameter (*D_N_*) in the coding regions has the value more than 325 repeats/Mbp which is three times greater than the *D_N_* value in untranslated regions (*D_N_* ∼ 95 repeats/Mbp) (Figure 17A). A similar observation has noticed in case of imperfect microsatellites also. *D_N_* values of imperfect microsatellites in coding and untranslated regions are 102 and 28 repeats/Mbp respectively (Figure 18A). Like microsatellites, minisatellites are found more in the coding regions than untranslated regions. *D_N_* values for perfect minisatellites in these regions are 24 and 14 repeats/Mbp respectively (Figure 17C) and in case of imperfect minisatellites, the values are 16 and 6 repeats/Mbp respectively (Figure 18C). While checking repeat density in length parameter (*D_L_*), perfect coding region repeats cover more than 0.3% of the genome in contrast to repeats in the untranslated regions which has the *D_L_* value of 0.09% (Figure 17B).

**Figure 17:**
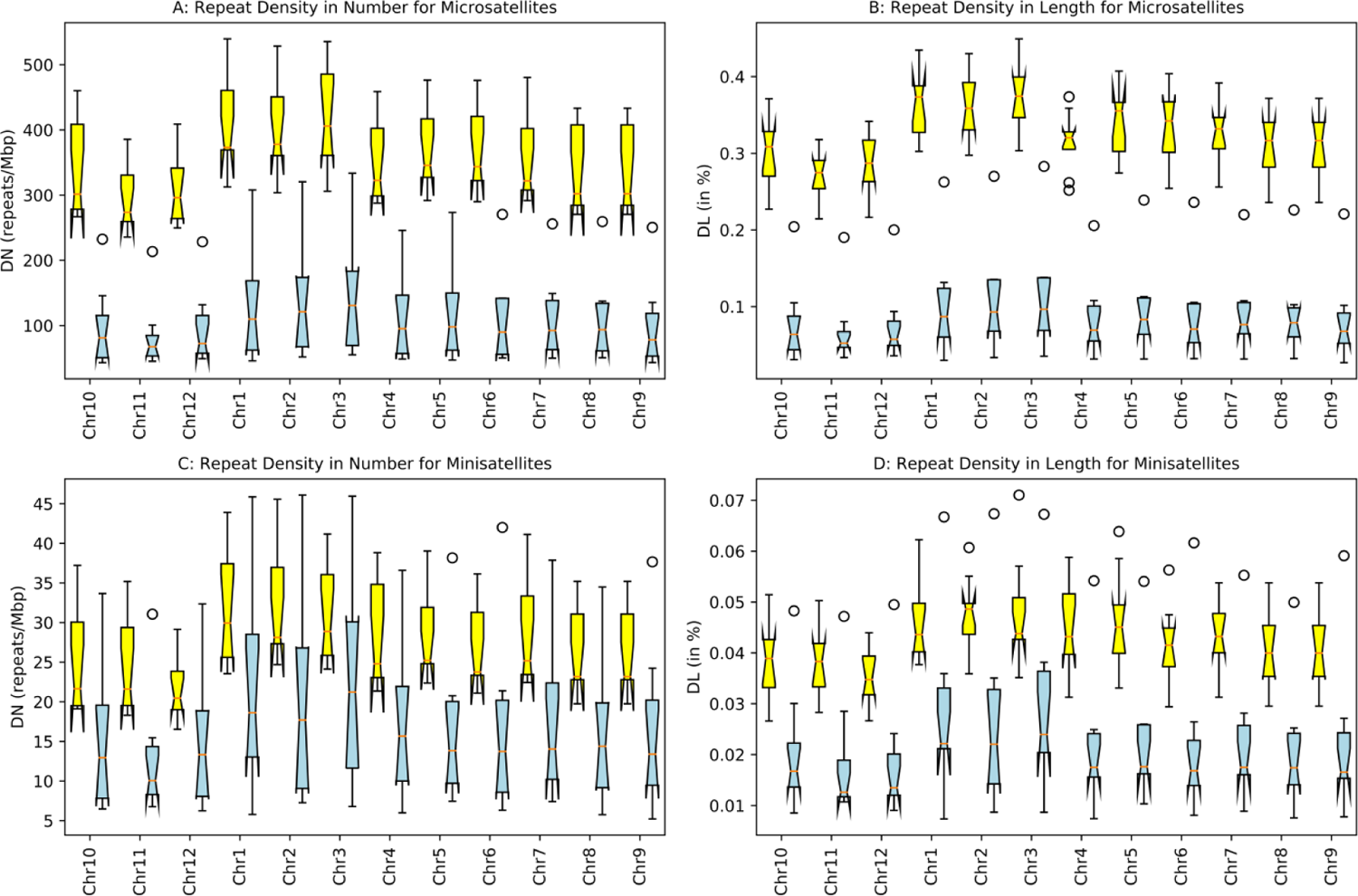
Chromosome-wise distribution of Perfect Tandem Repeats (Microsatellites and Minisatellites) in coding (Yellow) and untranslated regions (Blue).

**Figure 18:**
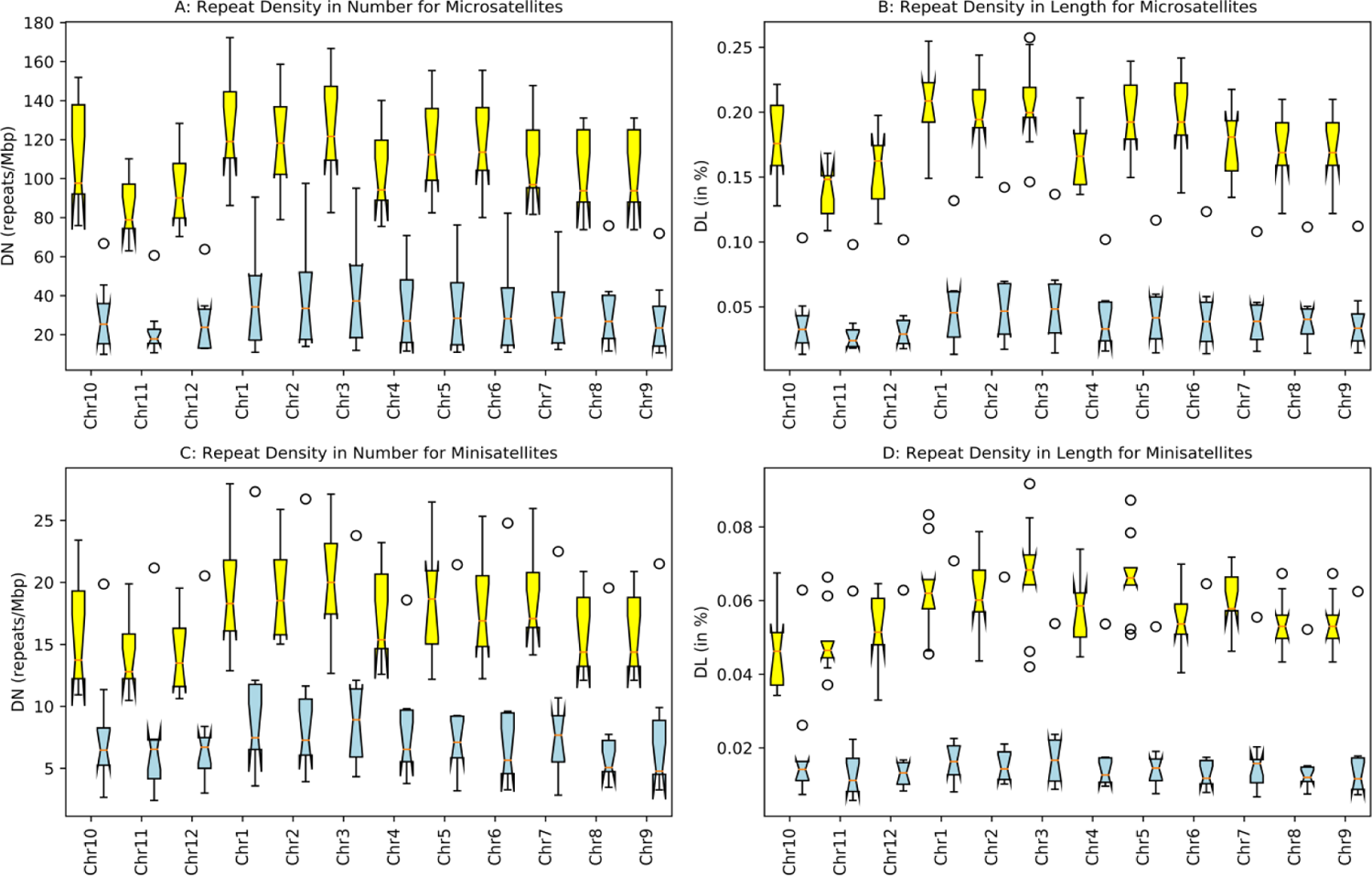
Chromosome-wise distribution of Imperfect Tandem Repeats (Microsatellites and Minisatellites) in coding (Yellow) and untranslated regions (Blue).

Imperfect microsatellites occur very less in the untranslated region (*D_L_* ∼ 0.04%) compare to coding region (*D_L_* ∼ 0.18%) (Figure18B). Perfect minisatellites have been detected slightly more in the coding regions (*D_L_* ∼ 0.04) compare to untranslated parts of the genome (*D_L_* ∼ 0.02%) (Figure 17D). *D_L_* value for imperfect minisatellites is moderately higher than perfect minisatellites (*D_L_* ∼ 0.06%) (Figure 18D). In comparison to coding regions, imperfect minisatellites in the untranslated regions cover only 0.02% of the genome. Regarding tandem repeats in coding and untranslated regions, *O. nivara* and *O. sativa* has the maximum number of perfect microsatellites in coding regions and untranslated regions respectively. One major observation is that the numbers of perfect microsatellites per Mbp in coding and untranslated regions in *O. sativa* genome are close to each other (*D_N_* ∼ 321 and 270 repeats/Mbp respectively) compare to the other species. In case of other genomes, as expected coding regions have more perfect microsatellites than untranslated region. Another prime observation is that the number of perfect minisatellites has occurred more in number in the untranslated regions than in the coding regions in case of *O. sativa* genome which might have some functional importance.

### Applications: Analysis of Common repeats

Analysis of common repeats has been performed using several groups. Figure 19A, 21A and 23A has shown clustered the species from Asia and 19B, 21B and 23B for the Africa continent using perfect, imperfect and interspersed repeats respectively. America has only one species hence continent wise grouping cannot be performed. Analysis shows presence of many species specific repeats across both Asian and African varieties. When comparisons between wild and cultivated species from three different continents (Asia, Africa and America has been done using perfect (Figure 20A), imperfect (Figure 22A), and interspersed repeats (Figure 24A), diversification of rice species has been observed. Similar observations have been noticed while comparing different genome types from aforementioned continents using perfect (Figure 5.20B), imperfect (Figure 22B), and interspersed repeats (Figure 22B). All of these results supports the rapid diversification of rice with time [36, 69]. To get more transparency on the results, pair-wise comparisons have been done using perfect (Figure 25) and imperfect repeats (supplementary Figure F2). Result has explained the origins of the cultivated species from their wild ancestors and shows similarity among species with same genome type. Anothet important observation is the grouping of species those belong to same continent. All these results indicates role of the repeats in evolution and their usability in ancestry prediction.

**Figure 19:**
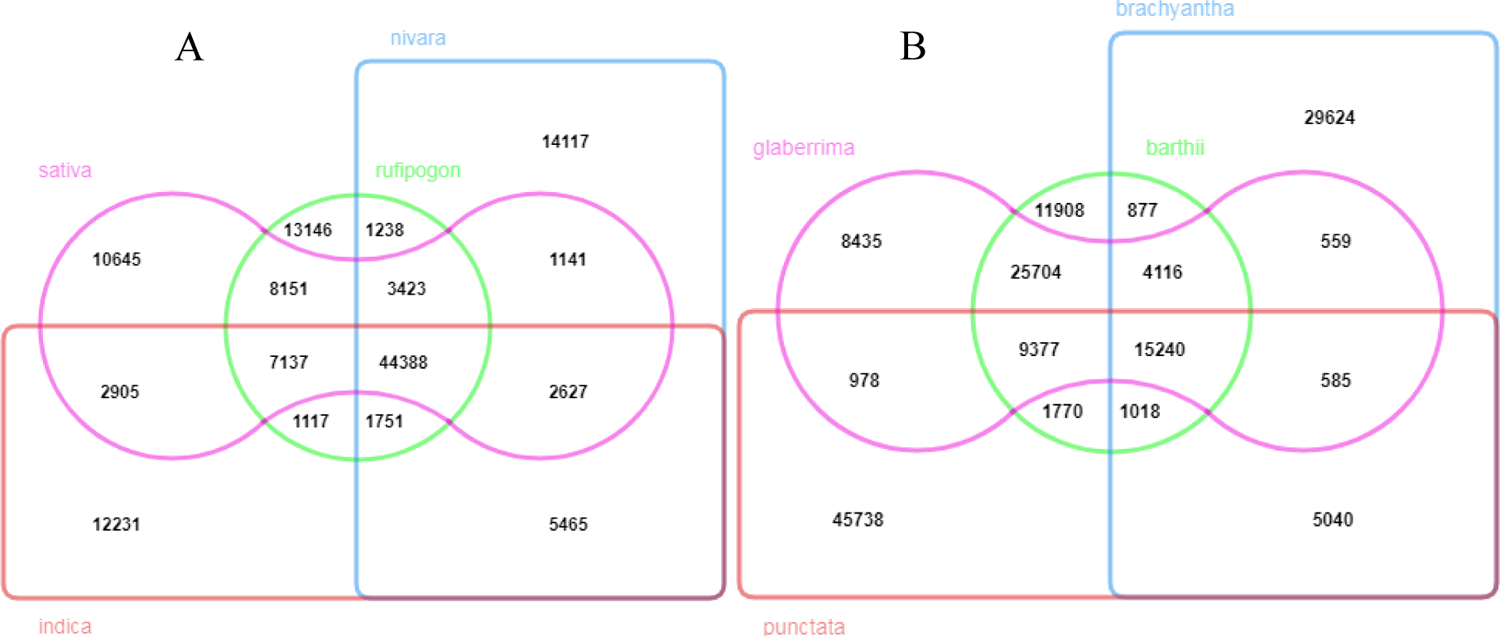
Venn diagram showing common perfect repeats in different species from A: Asia; B: Africa. Pink: *O. sativa*, *O. glaberrima*; Blue: *O. nivara*, *O. brachyantha*; Green: *O. rufipogon*, *O. barthii*; Red: *O. indica*, *O. punctata*.

**Figure 20:**
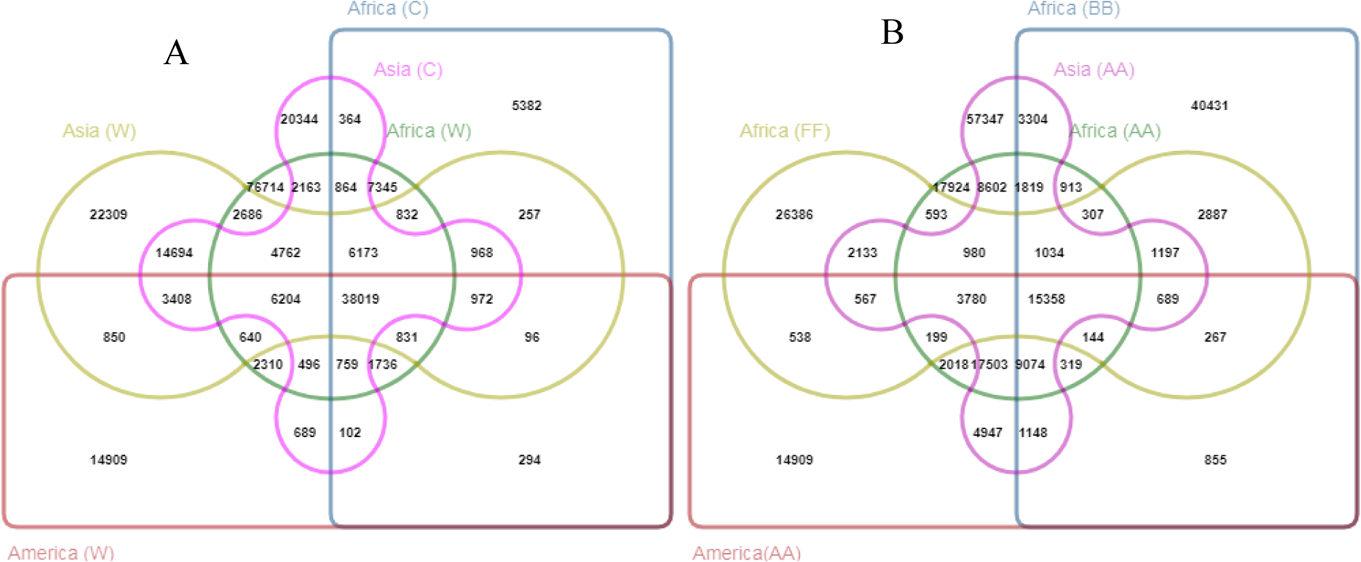
Venn diagram showing common perfect repeats in A: Wild and cultivated species from the different continents; B: Species from the different continents with different genome types. Pink: Asia (Cultivated), Asia (AA); Blue: Africa (Cultivated), Africa (BB); Green: Africa (Wild), Africa (AA); Red: America (Wild), America (AA); Yellow: Asia (Wild), Africa (FF).

**Figure 21:**
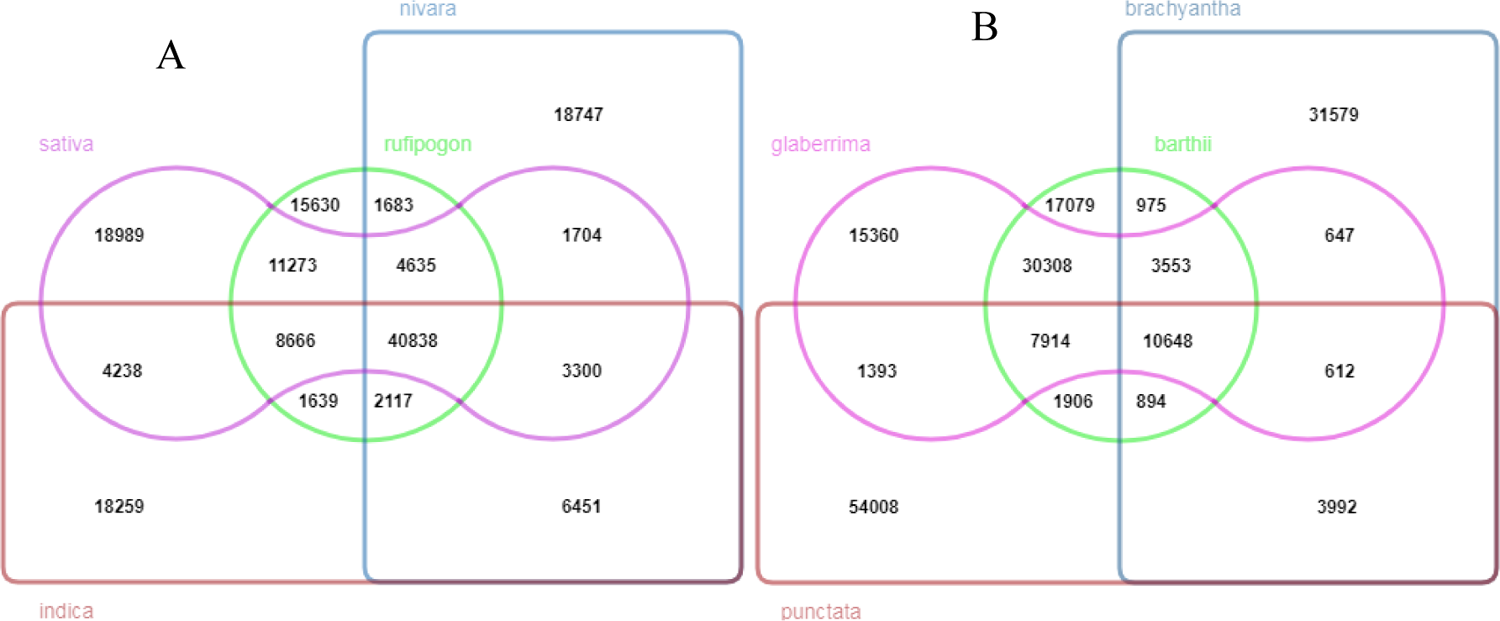
Venn diagram showing common imperfect repeats in different species from A: Asia; B: Africa. Pink: *O. sativa, O. glaberrima*; Blue: *O. nivara, O. brachyantha*; Green: *O. rufipogon, O. barthii;* Red: *O. indica, O. punctata*.

**Figure 22:**
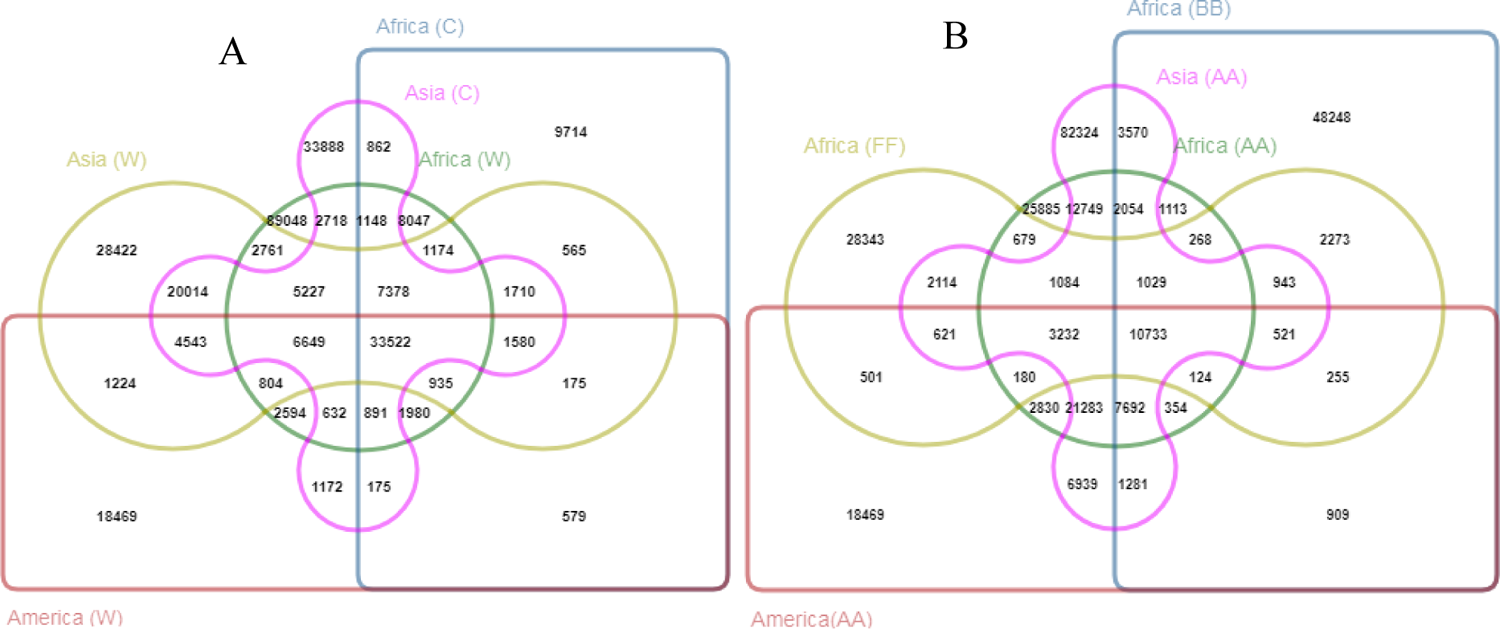
Venn diagram showing common imperfect repeats in A: Wild and cultivated species from the different continents; B: Species from the different continents with different genome types. Pink: Asia (Cultivated), Asia (AA); Blue: Africa (Cultivated), Africa (BB); Green: Africa (Wild), Africa (AA); Red: America (Wild), America (AA); Yellow: Asia (Wild), Africa (FF).

**Figure 23:**
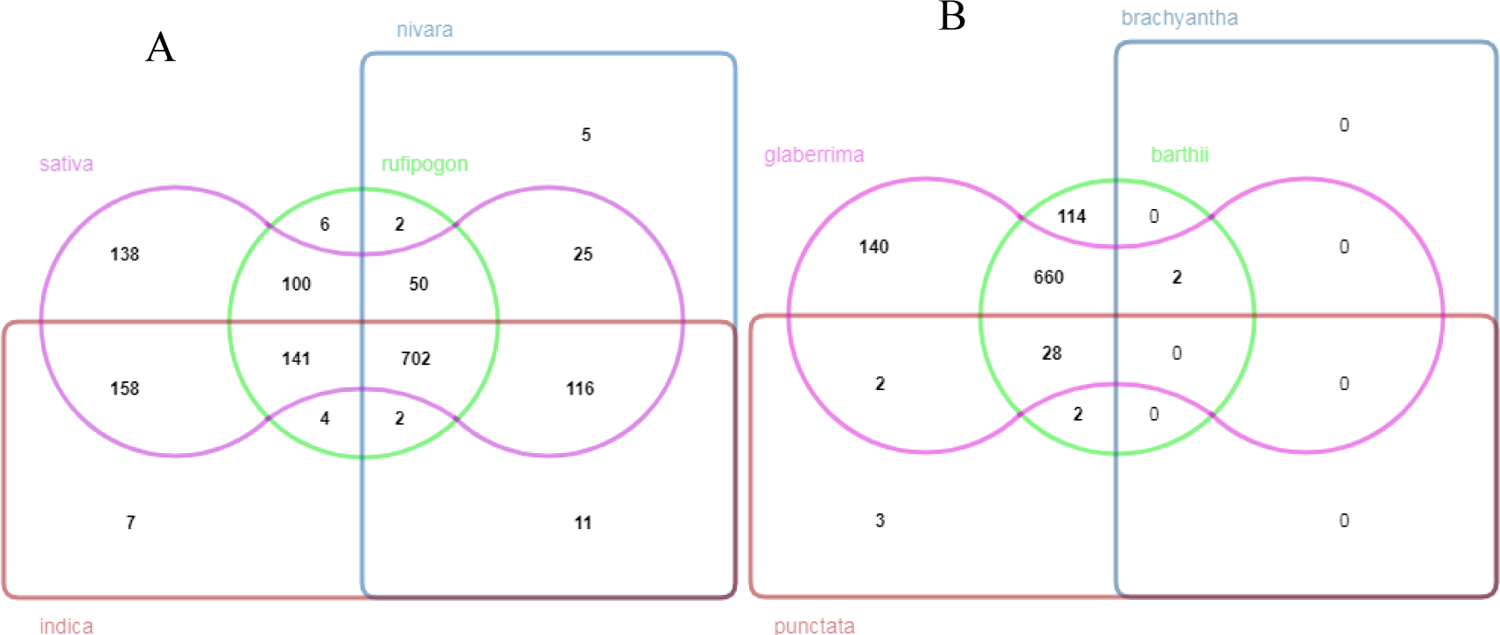
Venn diagram showing common interspersed repeats in different species from A: Asia; B: Africa. Pink*: O. sativa, O. glaberrima;* Blue*: O. nivara, O. brachyantha;* Green*: O. rufipogon, O. barthii;* Red: *O. indica, O. punctata*.

**Figure 24:**
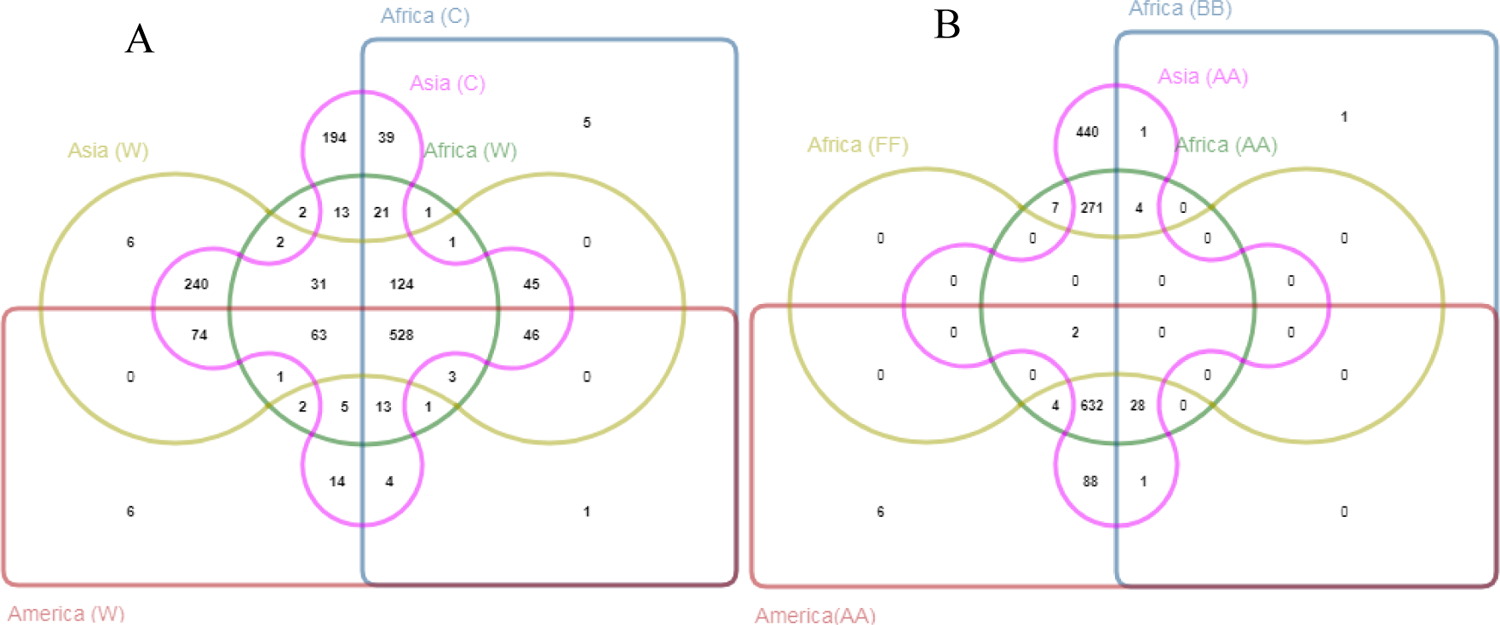
Venn diagram showing common interspersed repeats in A: Wild and cultivated species from the different continents; B: Species from the different continents with different genome types. Pink: Asia (Cultivated), Asia (AA); Blue: Africa (Cultivated), Africa (BB); Green: Africa (Wild), Africa(AA); Red: America (Wild), America (AA); Yellow: Asia (Wild), Africa (FF).

**Figure 25:**
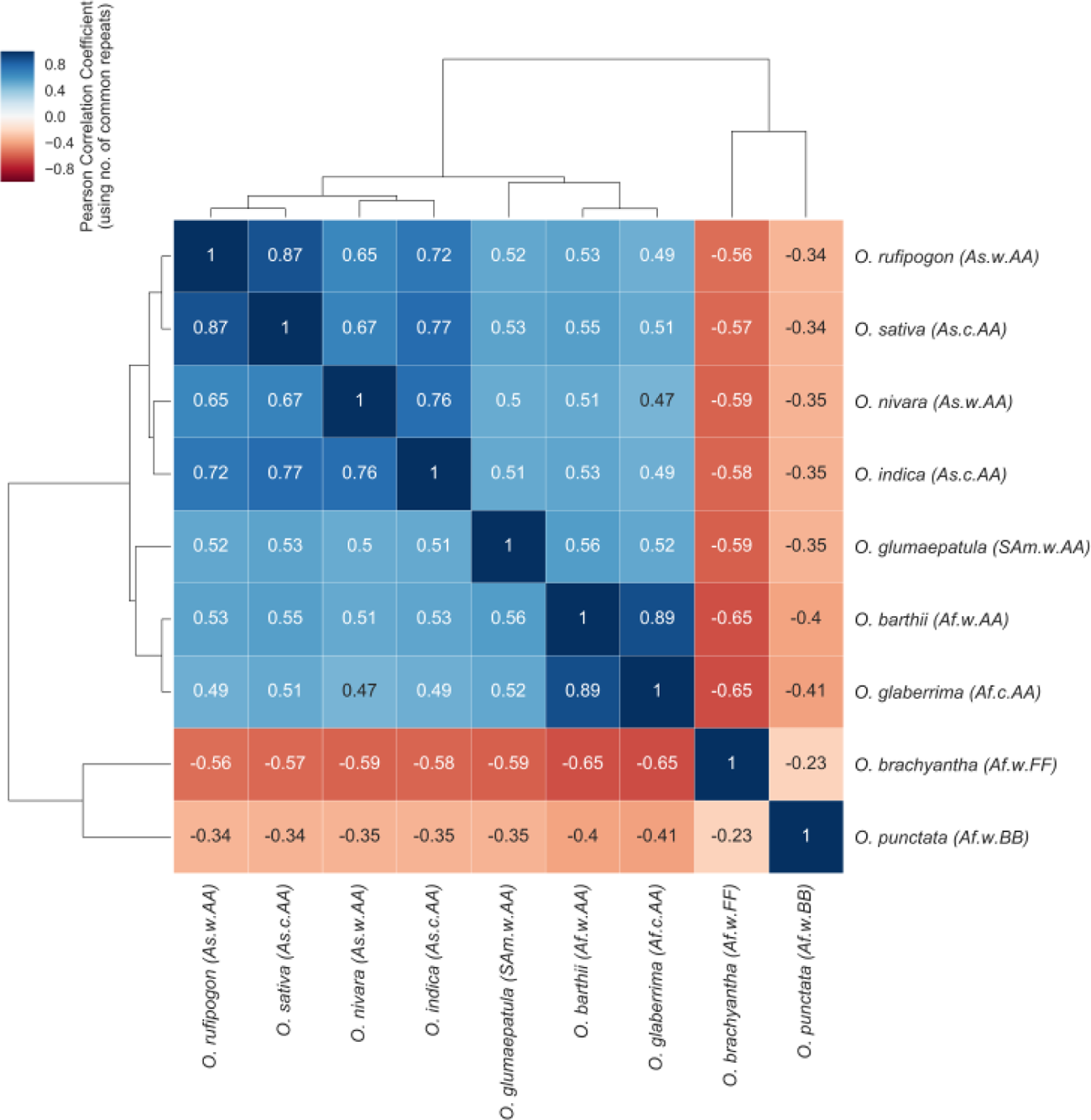
Pair-wise comparisons clustering of species using perfect common repeats. (Imperfect repeat: supplementary Figure F2). Red to Blue: Low to High Pearson correlation value; As: Asia, Af: Africa, Sam: South America; w: Wild, c: Cultivated.

### Correlation among genomic parameters and repeat densities

As shown in Table 4, repeat densities have no significant correlations with genomic parameters like Size (in Mbp), %GC content, %Genic regions and gene densities per kbp. Similar observations have been found in published literature also [70]. Only significant and strong positive correlation has been found between two repeat density parameters i.e. repeat density in length (*D_L_*) and repeat density in number (*D_N_*) (bolded in Table 4).

**Table 4:** Correlation Coefficients among Repeat Densities and genomic parameters

| Repeat Density | Genomic Parameter | Pearson | P-value* | Spearman | P-value* | Kendall | P-value* |
| --- | --- | --- | --- | --- | --- | --- | --- |
| **$D_L$** | Size(Mbp) | 0.0540 | 1.000 | 0.0333 | 1.000 | 0.0555 | 1.000 |
|  | %GC Content | 0.1981 | 1.000 | 0.2333 | 1.000 | 0.2777 | 1.000 |
|  | Gene Density per kbp | 0.8231 | 0.172 | 0.7999 | 0.260 | 0.6666 | 0.332 |
|  | Genic region (in %) | -0.2917 | 1.000 | -0.1166 | 1.000 | 0.0555 | 1.000 |
| **$D_N$** | **$D_L$** | **0.9528** | **0.00193** | **0.8833** | **0.043** | **0.7777** | 0.094 |
|  | Size(Mbp) | -0.0331 | 1.000 | -0.1499 | 1.000 | -0.0555 | 1.000 |
|  | %GC Content | 0.1737 | 1.000 | 0.0166 | 1.000 | 0.0555 | 1.000 |
|  | Gene Density per kbp | 0.7940 | 0.286 | 0.7333 | 0.662 | 0.5555 | 1.000 |
|  | Genic region (in %) | -0.1441 | 1.000 | 0.0333 | 1.000 | 0.0555 | 1.000 |

\*All p-values are adjusted using Bonferroni correction [71]

### Comparison of rice with other species

*Arabidopsis thaliana* and *Brachypodium distachyon* are the two species that have been chosen here for inter-species comparison. *A. thaliana* is the first model plant species which was sequenced in the year of 2000 due to its small genome size and short life cycle of about 6-8 weeks from its germination to mature seeds [72]. It is well studied and widely used in comparative genomics. On the other hand, *B. distachyon*is the wild monocot grass species that belongs to the same family as rice i.e. Poaceae. Like *Arabidopsis*, it has also small genome, short life cycle and can grow easily. These attributes have made it a suitable new model species for grass family [73].

### Comparing genomic parameters

As shown in the Figure 26, *Arabidopsis* has the lowest genome size of 119 Mbp followed by wild African rice species *O. brachyantha* and *Brachypodium*. %GC content is almost comparable in all of the species except *Arabidopsis* which has little less value of 36 Mbp (Figure 26A-26B). Though *Arabidopsis* has the smallest genome size among these species, but it has the maximum number of genes per kbp, highest amounts of %genic, %exonic and %coding region (Figure 26C-26F). Three cultivated rice species namely *O. glaberrima*, *O. indica,*and *O. sativa* have comparable gene density which is higher than the other rice species and *Brachypodium* but lower than *Arabidopsis.* Comparisons of the %genic, %exonic and %coding region have revealed that these values are comparable across all of the species except *Arabidopsis*. The observations regarding differences in genomic parameters in rice, *Arabidopsis,* and *Brachypodium* also justified because rice and *Brachypodium* belong to same monocot family whereas *Arabidopsis* which is a dicot belongs to different Brassicaceae (mustard) family.

**Figure 26:**
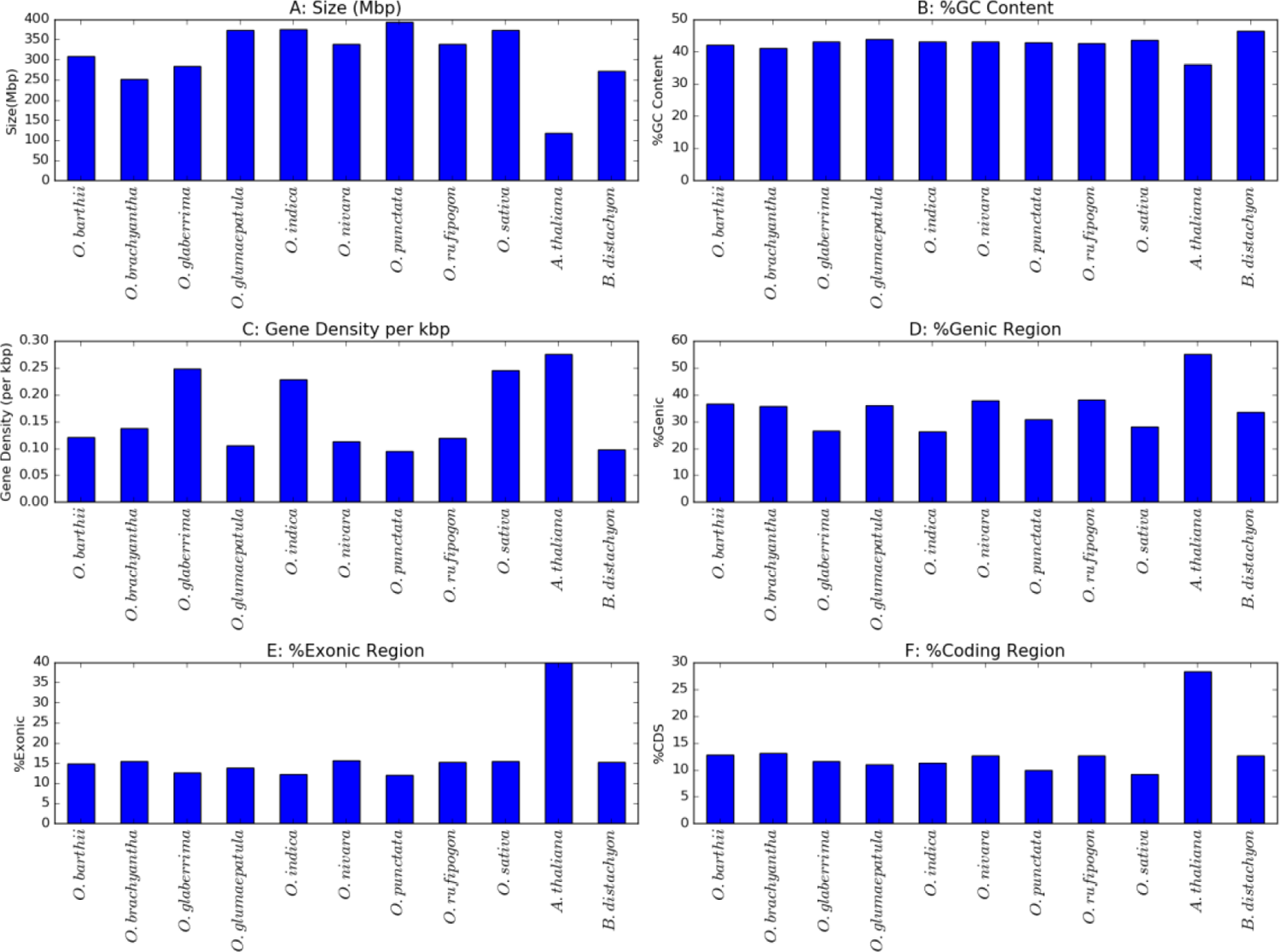
Comparison of genomic parameters across rice and other closely related species. A: Size in Mbp; B: %GC content; C: Gene density per kbp; D: % Genic region; E: % Exonic region; F: %Coding region

### Comparing repeat distributions

While comparing tandem and interspersed repeats (Figure 27) among the aforementioned plant species, the very first observation is that tandem repeats have been highly varied across all the species. In terms of repeat density in number parameter (*D_N_*), *O. sativa* has the highest value of 4182 repeats/Mbp whereas *B. distachyon* possesses the lowest value of 2770 repeats/Mbp. The median value is 3837 repeats/Mbp. In case of repeat density in length parameter (*D_L_*) also, *B. distachyon* contains the lowest amount of tandem repeats covering 3.7% of the genome in contrast to *O. sativa* genome which has approximately 6% of the genome covered by tandem repeats. The mean value for *D_L_* parameter is 5.07% with variance 0.4 which indicate that distribution of tandem repeats are highly varying across these plants (Figure 27A-27B).

**Figure 27:**
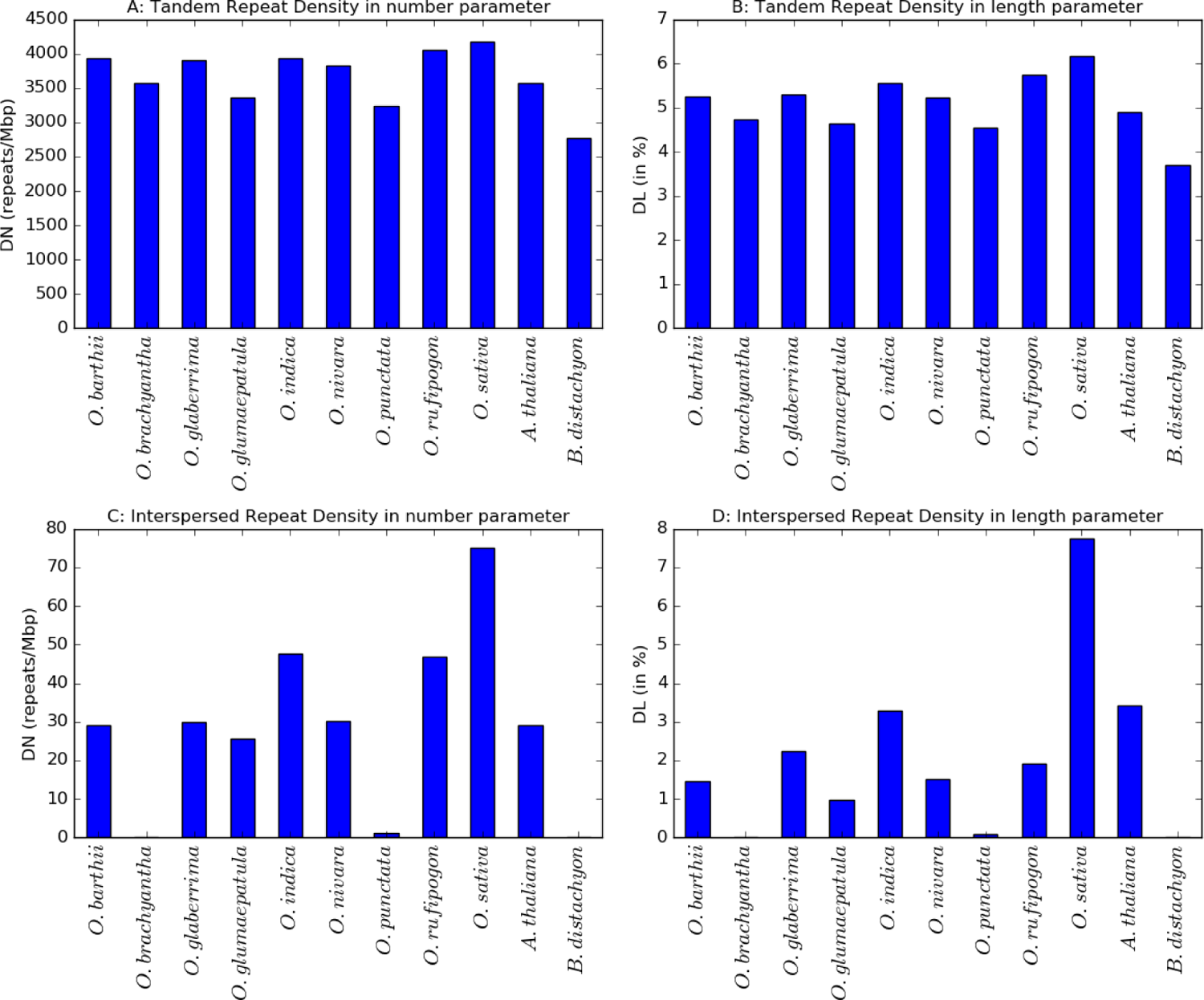
Comparing repeat distribution in *Oryza, Brachypodium,* and *Arabidopsis*; A: Tandem repeat density in number (*D_N_*); B: Tandem repeat density in length (*D_L_*); C: Interspersed repeat density in number (*D_N_*); D: Interspersed repeat density in length (*D_L_*);

Comparison of interspersed repeats has been performed too (Figure 27C-27D). In the Repbase v20.05 database, no entries have been found for *B. distachyon* species hence not included in the analysis. Comparing repeat density in number parameter has revealed that *O. sativa*has the highest number of interspersed repeats per Mbp (*D_N_* ∼75 repeats/Mbp) whereas *O. brachyantha* has the lowest number (< 1 repeat/Mbp). *Arabidopsis* has 29 repeats/Mbp in the genome, which is the median of *D_N_* values also. A similar observation has been found in case of repeat density in length parameter (*D_L_*). *O. sativa* has the highest coverage of 7.75% followed by *Arabidopsis* (*D_L_* ∼ 3.4%). *O. brachyantha* has the lowest value of 0.00072%. The median value for *D_L_* is 1.72%.

### Occurrences of repeats in stress-related genes

As mentioned earlier, sequencing most of the rice species have not been completed yet and annotations are in progress. Only for model species, *O. sativa* complete annotations have been provided in the RAP database [74]. Consequently, identification of stress-related genes has been performed mainly in the *O. sativa* species. Though, it is possible to detect stress-related genes in the other rice species using homology-based method utilizing *O. sativa* stress-related genes. But this may incorporate unreliable prediction in the analysis. Hence, in the present study of distinguishing stress related repeats, only *O. sativa* genome is considered for further analysis.

Tandem repeats have ubiquitously occurred in the rice stress-related genes. Set ‘A’ dataset of experimentally validated stress associated genes comprises of 2692 genomic loci out of which more than 99% of the loci contain perfect repeats (number of loci 2688) (Table 5). Set ‘B’ dataset of predicted stress-related genes, perfect repeats have occurred in all of the genomic loci. Approximately, 98% of the genomic loci that are related to housekeeping genes (the negative set of stress in the present study) contain perfect repeats. Similar observations have been found in case of imperfect repeats also. Imperfect tandem repeats have occurred in more than 99% of the genomic loci in both dataset ‘A’ and ‘B’ whereas 98% of the loci in the negative set have imperfect repeats. Distribution of interspersed repeats in the stress-related genes is quite less than tandem repeats. 29% of Set ‘A’ loci and 91% of Set ‘B’ contains interspersed repeats whereas only 6% of the housekeeping genes contain interspersed repeats.

**Table 5:** Repeat Occurrences in stress and housekeeping genomic loci

| Repeat Class | Dataset | Genomic loci available | Stress/Negative related loci with repeats | Repeats | Exclusive Repeats |
| --- | --- | --- | --- | --- | --- |
| Perfect | A | 2692 | 2688 (99.85%) | 11397 | 7052 |
|  | B | 952 | 952 (100%) | 4789 | 2635 |
|  | HK | 3644 | 3582 (98.30%) | 14277 | 9280 |
| Imperfect | A | 2692 | 2679 (99.52%) | 11260 | 7484 |
|  | B | 952 | 949 (99.68%) | 4558 | 2600 |
|  | HK | 3644 | 3571 (98%) | 12906 | 8571 |
| Interspersed | A | 2692 | 794 (29.49%) | 1234 | 1175 |
|  | B | 952 | 871 (91.49%) | 355 | 327 |
|  | HK* | 3644 | 235 (6.45%) | 1065 | 1015 |

\*HK stands for housekeeping

As shown in Figure 28, short tandem repeats are omnipresent in both the positive sets of stress-related genes but long repeats (motif length > 10 bp) are very rare. Trimeric repeating motifs are more frequent than other motifs followed by mononucleotide repeats and dinucleotide repeats. Imperfect hexanucleotide repeats are also marked their occurrences in the stress-related genes. Minisatellites of motif length 7 bp are found to occur in excess than the other minisatellites. Minisatellites are found to be more polymorphic than microsatellites in the stress-related genes.

**Figure 28:**
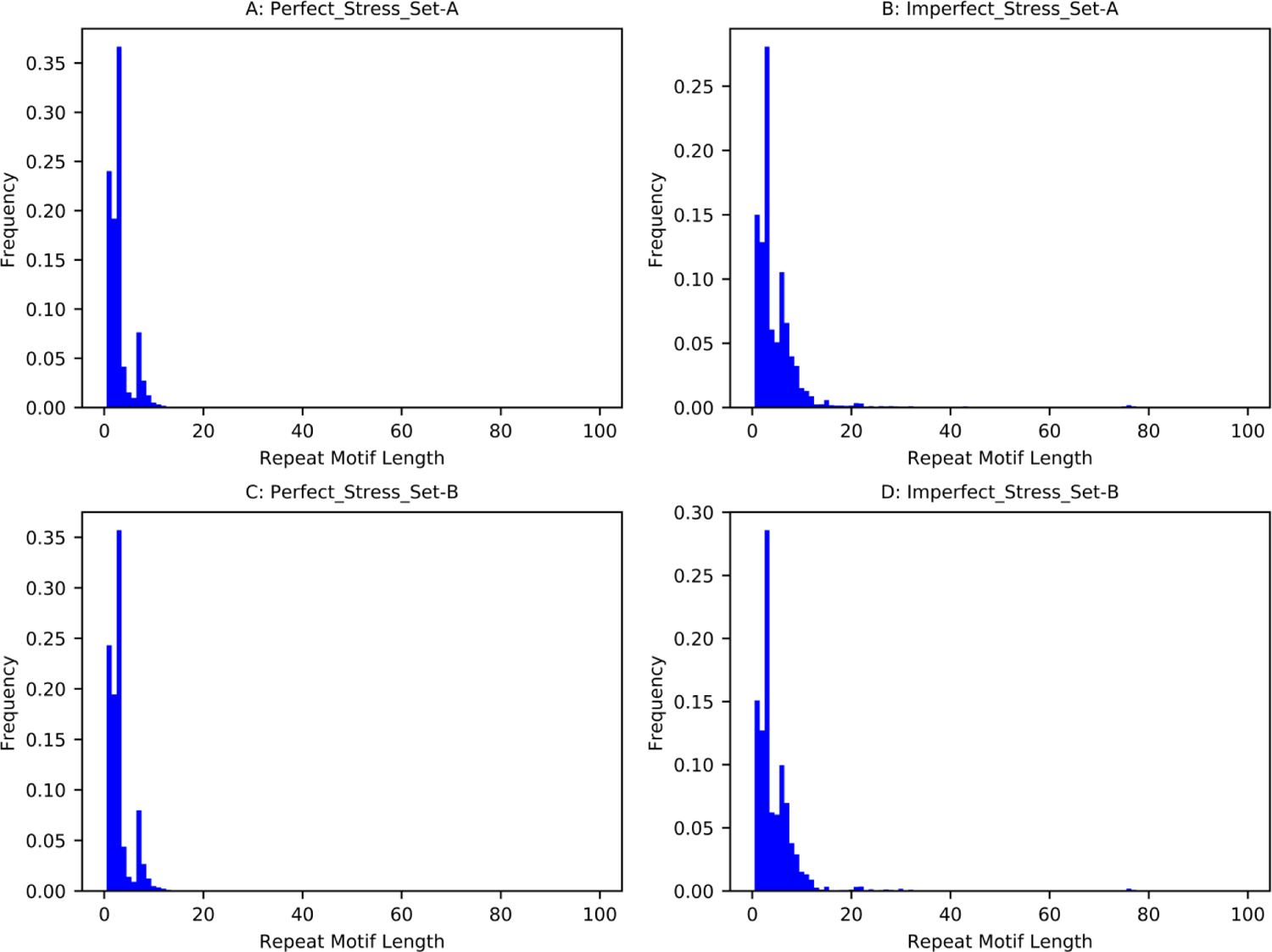
Distribution of repeats of different motif sizes; A: Perfect repeats in Set A gene set; B: Imperfect repeats in Set A gene set; C: Perfect repeats in Set B gene set; D: Imperfect repeats in Set B gene set;

While checking the number of loci that contain any repeat or not, it is found that repeats are not dispersed in the multiple genomic loci rather they are preferred to occur in particular locus (Figure 29). Similar observations have been found in case of both positive sets (A & B) and also valid for both perfect and imperfect repeats. Very few repeats have been found to occur in multiple genomic loci in rice as observed from Figure 29. This is the major difference that has been found between rice and *Salmonella.* In *Salmonella* repeats have occurred both in the single locus and in multiple loci. Present observation suggests that repeats in the rice genomes are highly precise and have some specific functional implications.

**Figure 29:**
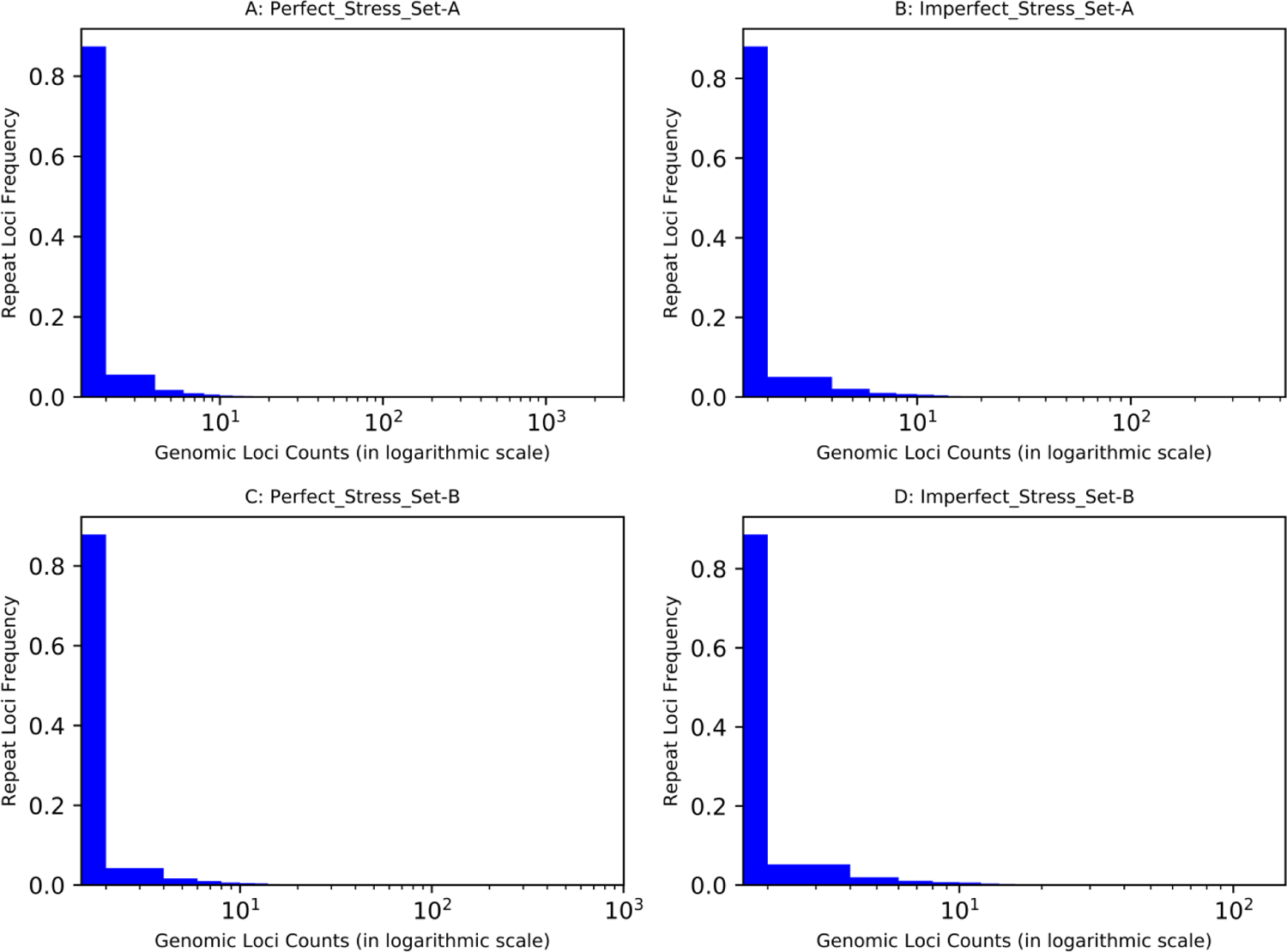
Distribution of repeats with varying genomic loci counts; A: Perfect repeats in stress genes (set A) B: Imperfect repeats in stress genes (set A); C: Perfect repeats in stress genes (set B); D: Imperfect repeats in stress genes (set B).

Distributions of repeats in stress-related and NSS genes (housekeeping genes) have been compared and whether there is an association between stress and repeat, have been tested using null hypothesis stated in Table 6. The comparison suggests that significant positive association has been observed between stress and repeats. Several tests including chi-square test for independence and Fisher exact test have been conducted to compare the occurrences of the repeats in the different genomic locations. Results have shown that repeat’s occurring in different stress-related genomic locations are significantly associated (p-value < 0.05) with stress-related genes. Similar observations have been found for all the repeat classes. To reduce Type I error in multiple comparisons of the genome, Bonferroni correction has been applied to adjust the p-values. Repeat classes, genomic locations, test statistics and corresponding p-values have been listed in Table 6.

**Table 6:** Comparison of repeats in stress and NSS genes

| **Null Hypothesis:** There is no association between repetitive elements and stress related loci |  |  |  |  |  |  |
| --- | --- | --- | --- | --- | --- | --- |
| Repeat Class | Location | Fisher's Exact test |  | Chi-square Test |  | Null Hypothesis Significance level 0.05 |
|  |  | Test Statistic | p-value* | Test Statistic | p-value* |  |
| Perfect Microsatellites | Inside gene | 15.74 | 3.30e-13 | 51.38 | 1.55e-11 | Rejected |
|  | Upstream | 15.73 | 3.32e-13 | 51.34 | 1.47e-11 | Rejected |
|  | Downstream | 15.75 | 3.32e-13 | 51.44 | 1.48e-11 | Rejected |
|  | All locations | 15.75 | 3.32e-13 | 51.44 | 4.52e-10 | Rejected |
| Perfect Minisatellites | Inside gene | 13.99 | 5.78e-12 | 44.73 | 1.20e-11 | Rejected |
|  | Upstream | 15.9 | 3.30e-13 | 51.84 | 1.07e-11 | Rejected |
|  | Downstream | 15.97 | 3.70e-13 | 52.06 | 1.50e-11 | Rejected |
|  | All locations | 15.75 | 1.68e-13 | 51.4 | 2.52e-11 | Rejected |
| Imperfect Tandem | Inside gene | 15.49 | 8.62e-14 | 50.39 | 1.41e-11 | Rejected |
|  | Upstream | 15.79 | 3.32e-13 | 51.52 | 5.22e-12 | Rejected |
|  | Downstream | 16.33 | 4.70e-19 | 53.48 | 1.49e-11 | Rejected |
|  | All locations | 15.75 | 3.94e-13 | 51.42 | 2.04e-17 | Rejected |
| Interspersed | Inside gene | 27.1 | 3.34e-15 | 78.02 | 2.80e-11 | Rejected |
|  | Upstream | 16.17 | 3.90e-15 | 50.19 | 2.08e-13 | Rejected |
|  | Downstream | 19.32 | 1.72e-13 | 59.82 | 2.88e-13 | Rejected |
|  | All locations | 18.31 | 3.30e-13 | 59.18 | 1.45e-11 | Rejected |
| All Repeats | Inside gene | 15.76 | 1.73e-13 | 51.47 | 1.49e-11 | Rejected |
|  | Upstream | 15.75 | 3.32e-13 | 51.42 | 1.45e-11 | Rejected |
|  | Downstream | 15.76 | 3.34e-15 | 51.47 | 1.48e-11 | Rejected |
|  | All locations | 15.75 | 3.90e-15 | 51.44 | 2.88e-13 | Rejected |

\*All p-values are adjusted using Bonferroni correction [75]

### Mining associations between stress and repeats

As shown in Table 5, total 16186 perfect and 15818 imperfect tandem repeats (cumulative count of Set A and Set B) have been identified in the stress-related datasets (both Set ‘A’ and Set ‘B’). Total 1589 interspersed repeats have been also detected in the stress-related genes. Extracting exclusive repeats and removing redundancies result 9086 (∼ 56%) perfect and 9537 (∼ 60%) imperfect tandem repetitive loci. On the other hand, 1502 (∼ 94%) exclusive interspersed repeats have been discovered. 493 and 420 exclusive perfect and imperfect tandem repeats have been found to be common between Set ‘A’ and Set ‘B’ stress gene datasets. While finding common exclusive interspersed repeats, only 73 repetitive loci have been distinct. Total 19701 unique exclusive repetitive loci (combining perfect, imperfect and interspersed and removing redundancies) has been identified, out of which more than 99% (∼ 19631) have shown positive association (nPMI> 0) with stress (Figure 30). Lists of top 10 potential stress associated perfect, imperfect and interspersed repeats (sorted by nPMI score) have been presented in Table 7, Table 8 and Table 9 respectively and discussed below along with its product information. These repeats can be used as the markers for identifying novel stress-related genes in *Oryza* species.

**Figure 30:**
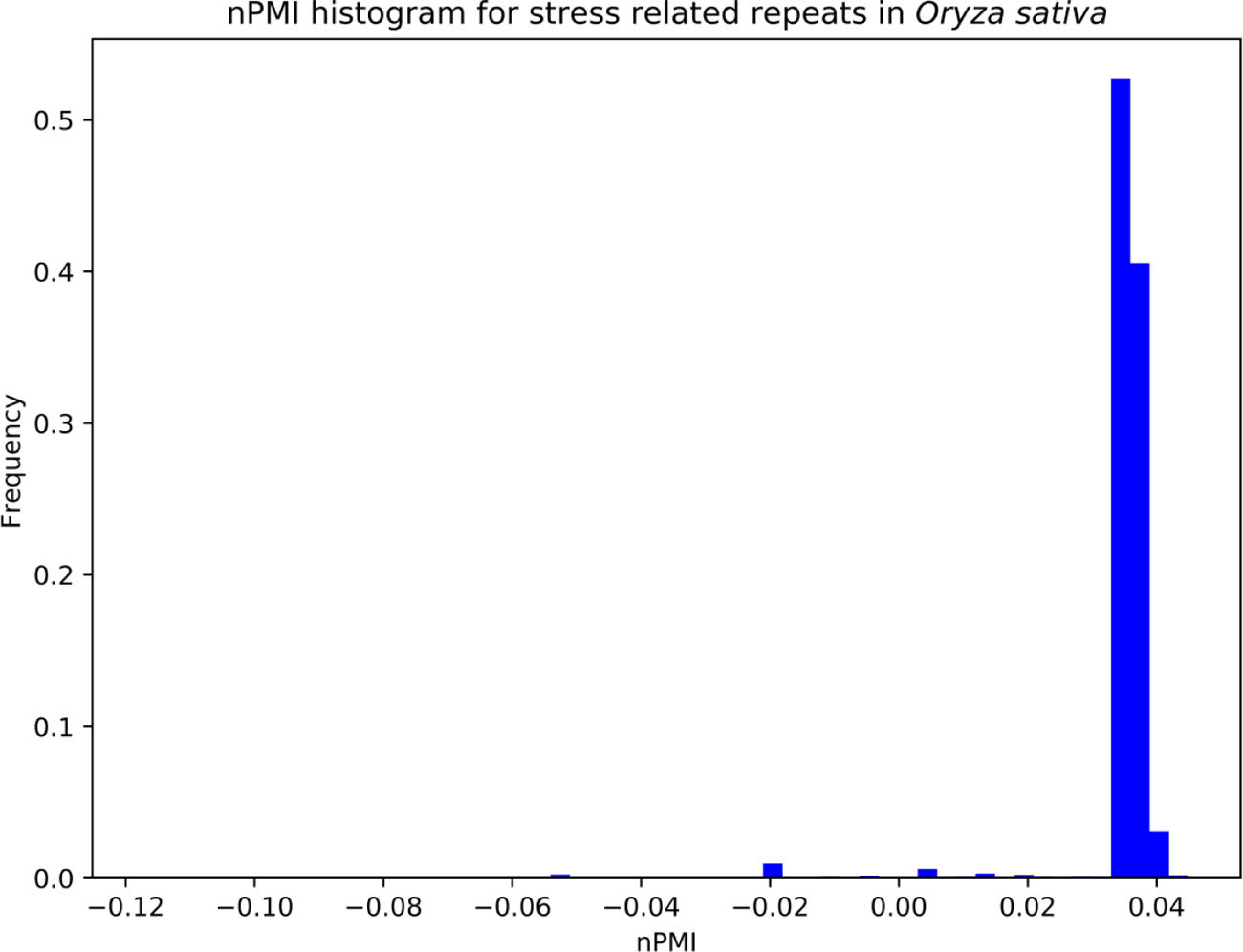
Distribution of nPMI values of exclusive stress related repeats in *O. sativa* species.

**Table 7:**
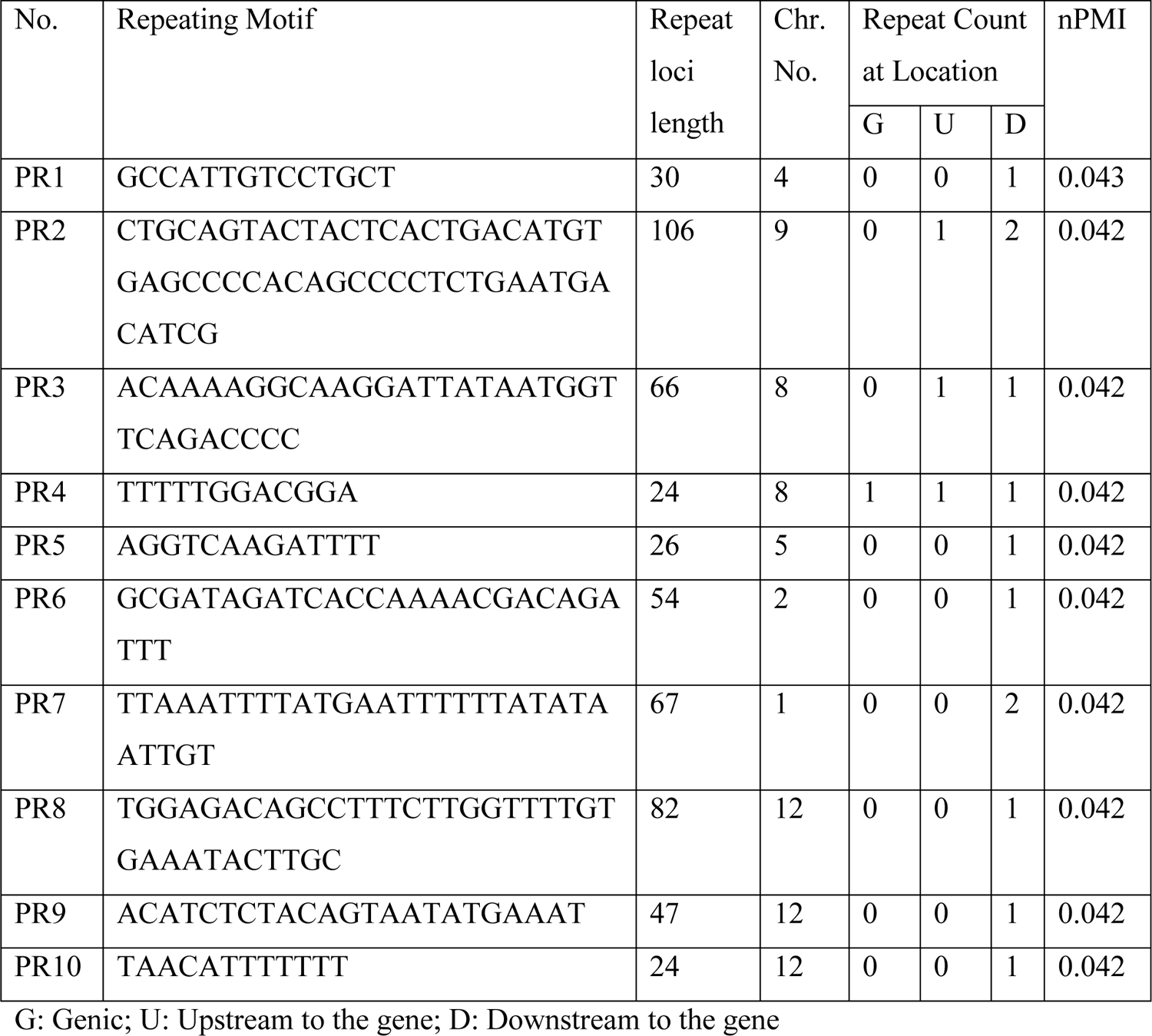
Exclusive Perfect Minisatellites (Motif length > 10) sorted by nPMI

**Table 8:**
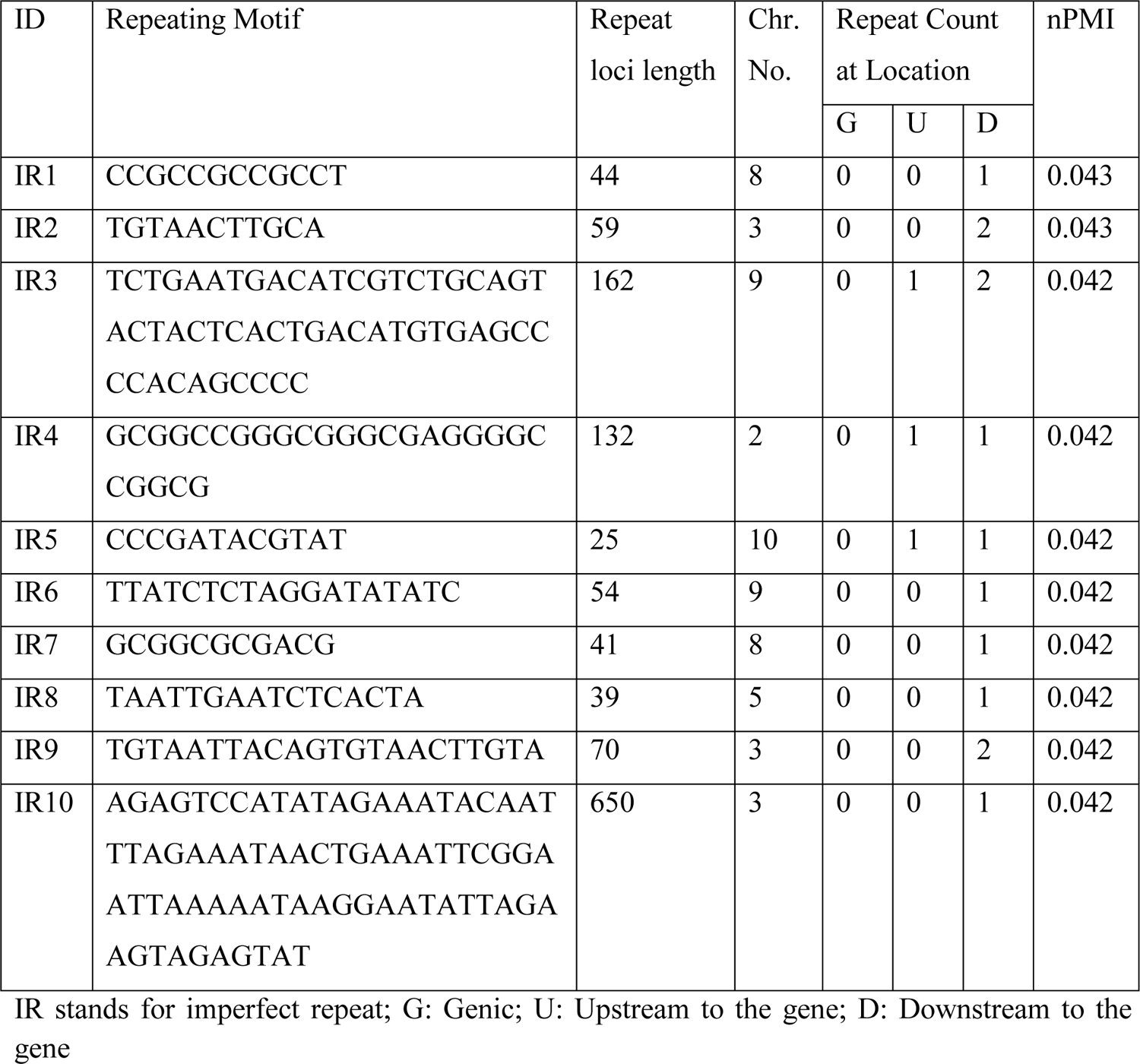
Exclusive Imperfect minisatellites (Motif length > 10) sorted by nPMI

**Table 9:**
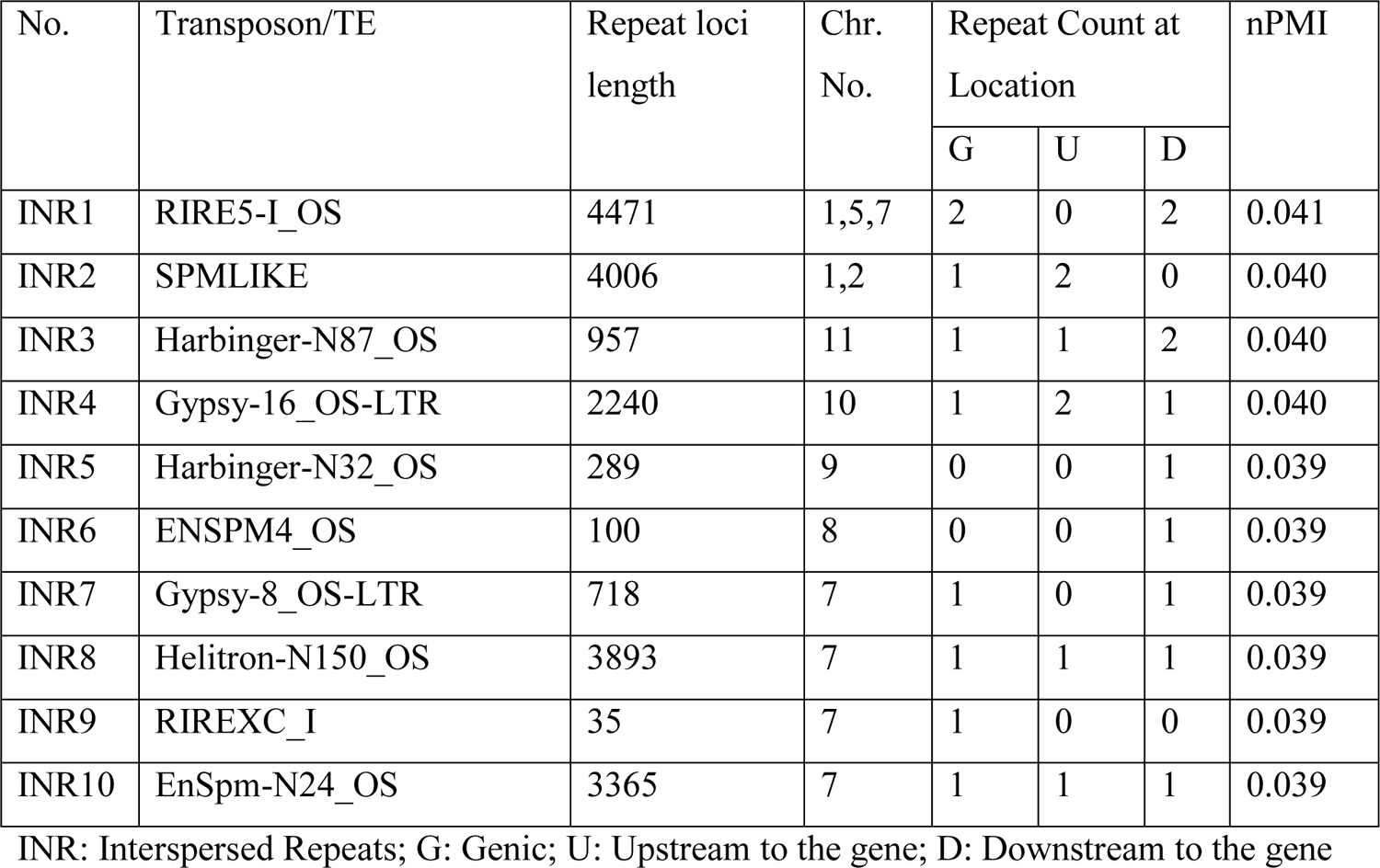
Interspersed repeats exclusively present inside or in vicinity of stress genes

| No. | Transposon/TE | Repeat loci length | Chr. No. | Repeat Count at Location |  |  | nPMI |
| --- | --- | --- | --- | --- | --- | --- | --- |
|  |  |  |  | G | U | D |  |
| INR1 | RIRE5-I_OS | 4471 | 1,5,7 | 2 | 0 | 2 | 0.041 |
| INR2 | SPMLIKE | 4006 | 1,2 | 1 | 2 | 0 | 0.040 |
| INR3 | Harbinger-N87_OS | 957 | 11 | 1 | 1 | 2 | 0.040 |
| INR4 | Gypsy-16_OS-LTR | 2240 | 10 | 1 | 2 | 1 | 0.040 |
| INR5 | Harbinger-N32_OS | 289 | 9 | 0 | 0 | 1 | 0.039 |
| INR6 | ENSPM4_OS | 100 | 8 | 0 | 0 | 1 | 0.039 |
| INR7 | Gypsy-8_OS-LTR | 718 | 7 | 1 | 0 | 1 | 0.039 |
| INR8 | Helitron-N150_OS | 3893 | 7 | 1 | 1 | 1 | 0.039 |
| INR9 | RIREXC_I | 35 | 7 | 1 | 0 | 0 | 0.039 |
| INR10 | EnSpm-N24_OS | 3365 | 7 | 1 | 1 | 1 | 0.039 |

INR: Interspersed Repeats; G: Genic; U: Upstream to the gene; D: Downstream to the gene

### Repeats occurring exclusively in stress-related genes of *Oryza sativa*

According to the present analysis of stress associated perfect repeats, approximately 89% of the repeats are found to be minisatellites those are exclusively present in the Set A (gold standard) stress-related genes. A similar result (∼90%) has been found for stress-related genes in the Set B also. Majority of these minisatellites have occurred in the upstream or downstream regions of the genome. Very few of these repeats have been found inside the gene.

As mentioned before, Table 7 shows top 10 (sorted by nPMI values) perfect minisatellites of length greater than 10 nucleotides those are exclusively associated with stress. 14-mer repeating motif ‘GCCATTGTCCTGCT’ with length 30 solely found in the *O. sativa* species in the locus that encodes for heat shock protein (HSP90). Heat shock proteins are a class of molecular chaperones those are crucial for plant growths and related to several important functions like cell signaling, cell immunity, integrity maintenance and many more [76].

Another repeat of motif length 53 bp is related to the DNA binding protein coding gene possibly a transcription factor (TF) (locus found in Grass TFDB [77]) of basic helix-loop-helix (bHLH) family. This family of TFs has role in phytochrome reaction under photoreceptor signaling pathway. Other occurrences of this motif are in the downstream region of carbonate dehydratase encoding gene and upstream of an unknown/hypothetical protein-coding gene. This particular repeat is also present in the *O. rufipogon* species which is thought to be the ancestor of *O. sativa* [36]. A 33-mer repeating motif “ACAAAAGGCAAGGATTATAATGGTTCAGACCCC” of length 66 bp is present in the upstream region of the gene having an annotation of CCT domain containing protein which controls the flowering time also has been reported to be involved in abiotic stress response [78].

The same repeat has been also located in the downstream region of the gene encoding Succinate dehydrogenase iron protein beta subunit (SDBH) which serves as the direct source of reactive oxygen species (ROS) production in rice are also accompanied with up-regulation of many stress-related genes [79]. A complete list of all the exclusive stress associated perfect repeats (both micro- and minisatellites) has been listed in the Supplementary Table file.

Polymorphic stress related minisatellites which are only present in stress genes (Set A and Set B) have been detected too. Top 10 have been shown in Table 8. Like perfect repeats, major proportions (∼ 65%) of the imperfect repeats are in the upstream and downstream regions of the stress-related genes. A 12-mer polymorphic repeat of length 44 bp has occurred in the downstream region of the ROS stress controlling gene SDB [79]. Another repeating motif “TGTAACTTGCA” with repeat length 59 has occurred in the downstream regions of two important proteins. One is the IQ calmodulin-binding region domain containing protein which is very much essential in elevation of controlling Ca^2+^ concentration during the cross-talk of various stressors like cold, heat, drought, salt etc. [80]. The other protein is LSTK-1 kinase is responsive to drought and osmosis stress [81]. “TCTGAATGACATCGTCTGCAGTACTACTCACTGACATGTGAGCCCCACAGCCCC” is another repeating motif of length 54 bp have been exclusively associated with bHLH family of TFs and carbonic dehydratase like proteins whose roles in stress response have been discussed earlier. A complete list of stress associated imperfect repeats has been presented in the Supplementary Table file.

Top 10 transposons like interspersed repeats predicted to be associated with stress response have been listed in Table 9. LTR retrotransposon RIRE5-I_OS have been found inside and downstream region of the two proteins namely non-cyanogenic beta-glucosidase and serine carboxypeptidase2 which are involved in plant defense against biotic factors and oxidative stress [82–83]. A DNA transposon SPMLIKE has been found in the upstream regions of glycoside hydrolase and nitric oxide synthase. The glycoside hydrolase family proteins has a functional role in forming cell wall architecture and often shows their responses under stressful conditions [84]. Nitric oxide synthase is responsible for the production of nitric oxide and reported to be associated with salt stress response in *Arabidopsis* [85]. List of all exclusively stress related interspersed repeats are given in the Supplementary Table file.

### Analysis of false positives

Many repetitive sequences are found to occur in both stress and NSS or negative sets of genes (housekeeping genes) and marked as the false positives. Total 4345 perfect (9% of the total repeats associated with stress and negative gene sets), 3776 imperfect (8%) and 689 interspersed (1%) repeats have been detected as false positives in gold standard gene set (Set A). Similarly, 2154 perfect (∼5%), 1958 imperfect (∼ 3.5%) and 175 interspersed (∼ 0.3%) repeats have been identified as false positives in Set B (silver standard) stress-related genes. Out of 4345 perfect false positives in Set A, 61% are found to be minisatellites and 24% of the 3776 imperfect minisatellites have been marked as false positives. Similarly, in the Set B gene list, 49% of 2154 perfect, 14% of 1958 imperfect minisatellites have been identified as false positives. As mentioned earlier, housekeeping genes have been used as the negative set of for comparing repeat distributions but authentic housekeeping genes are few in number. Here, the list of housekeeping genes used has been predicted from expression data sets. Hence, there exists a high possibility that the repeats marked as false positives might be related to stress response but due to data insufficiency, they are predicted as false positives (Supplementary Table file).

### Prediction of novel candidate stress associated loci in *Oryza* genomes

Those repeats which are exclusively occurred in stress-related genes of *O. sativa* genome can be used as a marker to identify new candidate stress associated loci from other rice species. Some of the repeating motifs those are extracted earlier (Table 7) have been used to mark stress related loci in the genomes of rice species other than *O. sativa* as shown in Table 10. A 53-mer perfect minisatellite is present in the vicinity of the gene that encodes for carbonate dehydratase-like protein and located on chromosome 9 in *O. sativa*. The same repeat has been also found in the vicinity of “ORUFI09G14140” locus in *O. rufipogon* located in the same chromosome no. 9. The length of the repetitive sequence is 106 bp which is very much unlikely to be a random event.

**Table 10:**
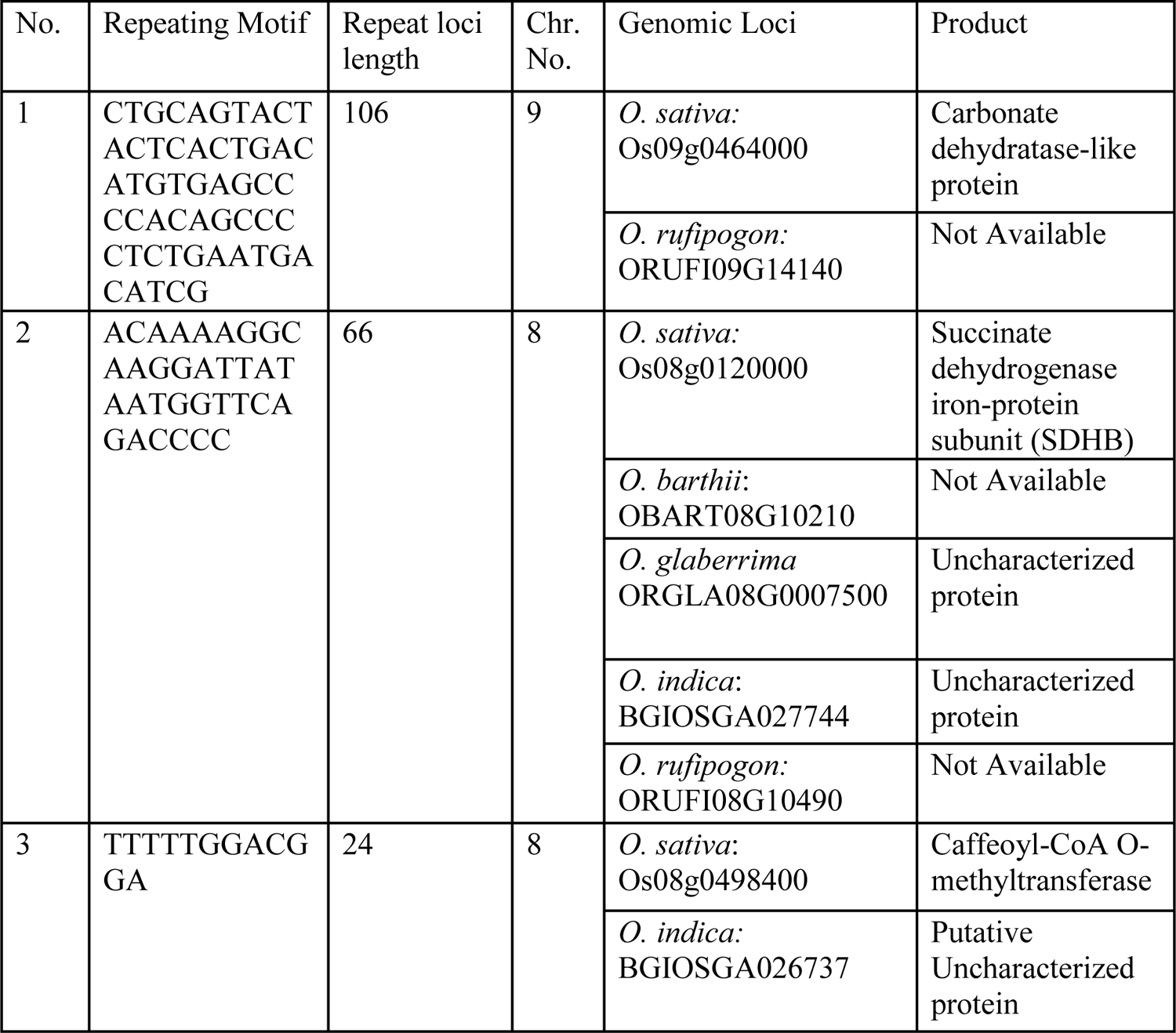
Prediction of novel candidate stress associated genes using repetitive markers

## Conclusions

Global temperature has been rising day by day due to the increment of the pollution level and emission of the greenhouse gases. As a result of this global warming accompanied with other stress factors, the severe adverse effect has been noticed on the rice yield in past few years. The exponential increment of the world’s population has magnified this negative impact up to several folds. To understand the stress response mechanisms for adaptation under unfavorable conditions and to increase the annual yield of rice to secure the upcoming food crisis, extensive studies are required. In the present study, a comparative genomics-based approach has been developed to detect and characterize repetitive sequences that might play an important role in adaptation under stressful conditions. Significant non-random distributions of repeats have been observed in rice. In case of tandem repeats, trimeric are found to be present in excess followed by monomeric and dimeric repeats. In comparison to short tandem repeats or microsatellites, long repeats (motif size > 10 bp) are less frequent but have been marked their occurrences in the genomes. Repeats are ubiquitously found in all genomic regions which include genic, intergenic, exons, introns and UTRs. No significant correlation has been observed among repeat densities and genomic parameters. Analysis of common repeats across 9 rice species has shown that each of species has some species-specific repeats. Most importantly, pairwise comparisons have clearly distinguished clusters of rice species from same geographical locations and of a particular genome type (Figure 23). Comparisons with other species like *Arabidopsis* and *Brachypodium* have strengthened this as similar results found from earlier published works.

Presence of repeats in known stress-related genes has been observed in the analysis. Repeat’s significant association with stress clearly indicates its functional role in adaptation under stressful environments. Though, experimental data insufficiency is a major obstacle to using it as a biomarker and to identify candidate stress related loci which are still un-annotated or hypothesized in rice species other than *O. sativa.* But, it can be a starting point to use repeats for novel gene identification of functional importance as done in the present work. However, it may not be used with certainty as a generic predictive method as repeats are occurring in almost every important known locations of the stress associated genes and genomes, so experimental validation is required along with the understanding its causal relation with stress response and adaptation.

