## Supplementary Figure for "Exploring repeats in rice genomes: Identification, Characterization and its Applications"

### Appendix: Chapter5

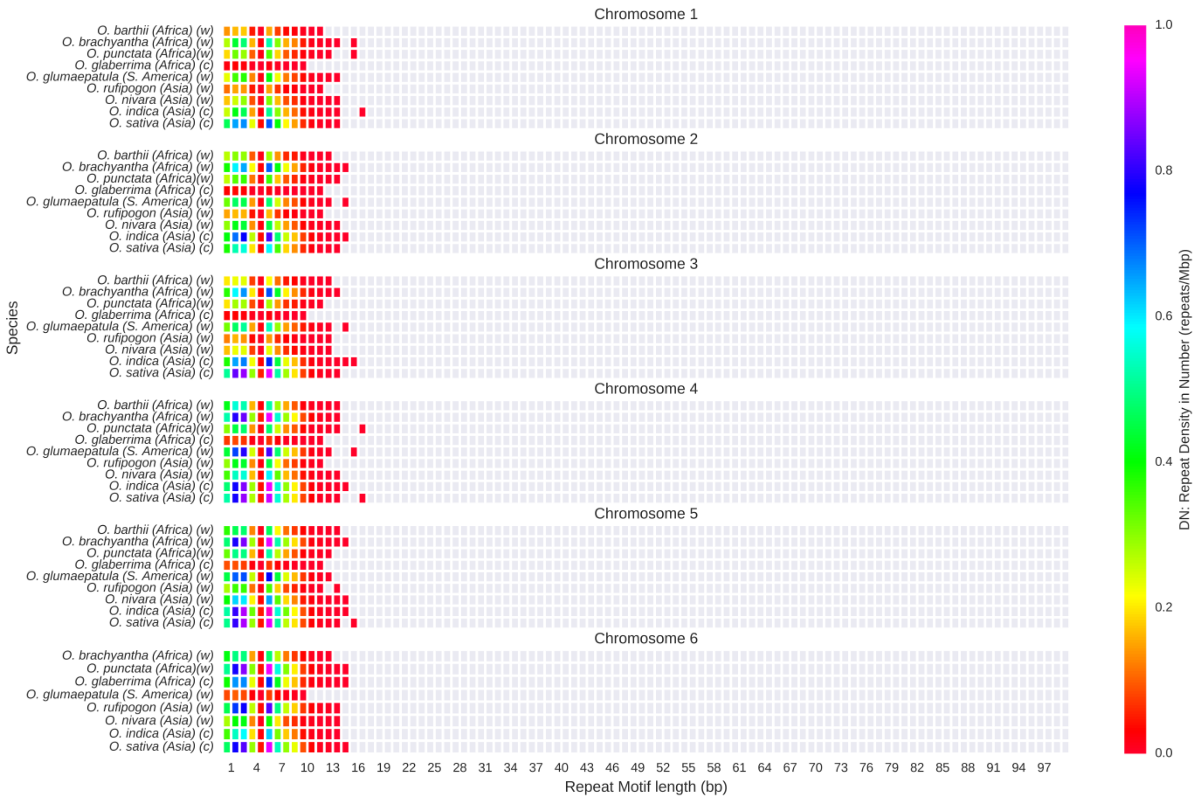


Figure A5.1A: Distribution of imperfect repeats in random sequences of rice chromosomes 1-6.

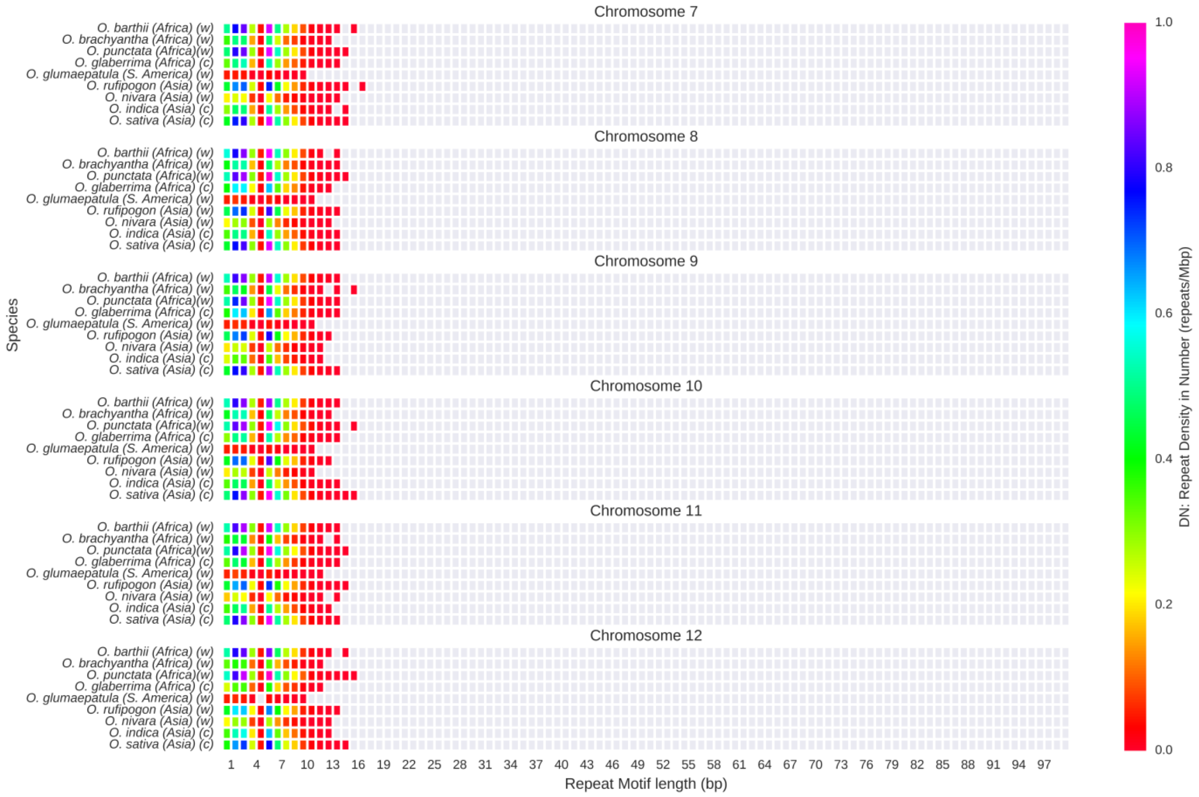


Figure A5.1B: Distribution of imperfect repeats in random sequences of rice chromosomes 7-12.

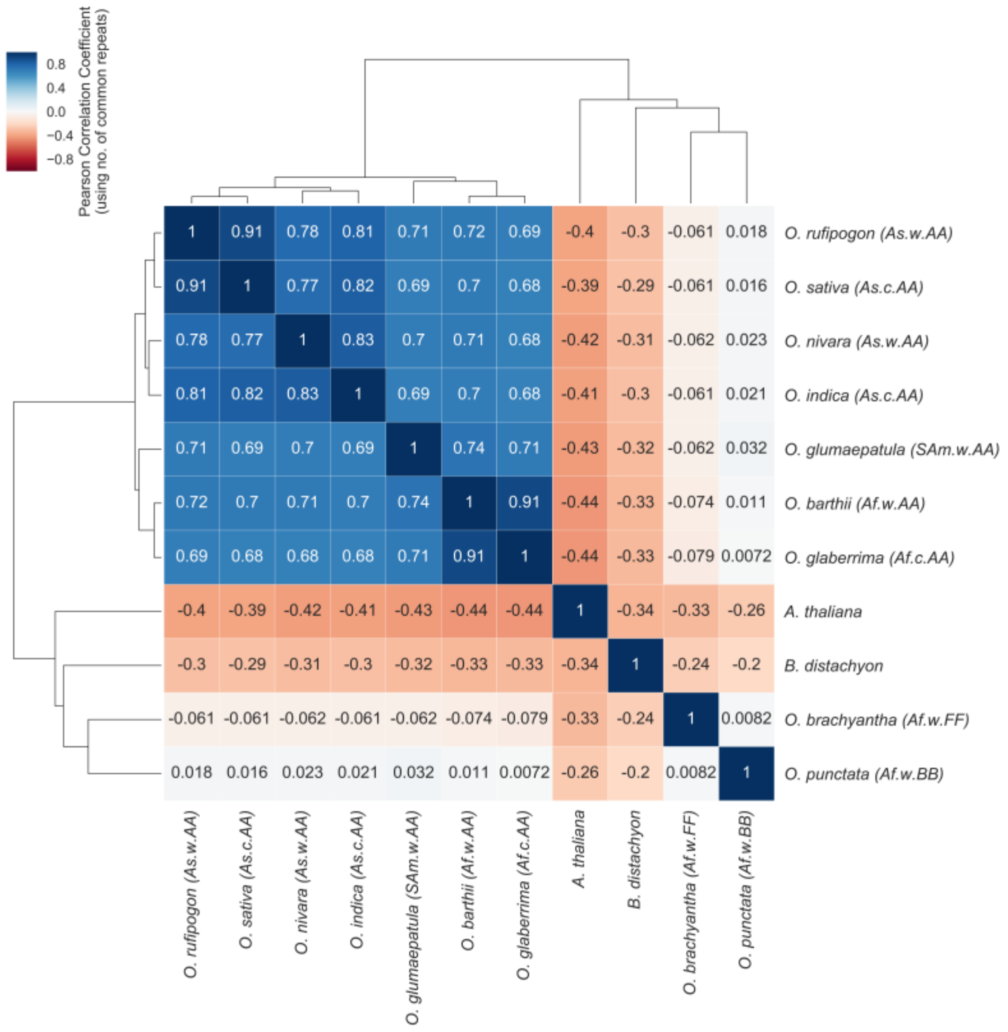


Figure A5.2: Pairwise comparisons and clustering of species using imperfect common repeats. Red to Blue: Low to High Pearson correlation value; As: Asia, Af: Africa, Sam: South America; w: Wild, c: Cultivated.
